# TLR4 signaling in presence of HIV-induced activation enhances programmed death ligand-1 expression on human plasmacytoid dendritic cells and modulates their function

**DOI:** 10.1101/2022.07.05.498853

**Authors:** Meher Patel, Sukhwinder Singh, Amy Davidow, Jihong Dai, Patricia Fitzgerald-Bocarsly

## Abstract

Human Plasmacytoid dendritic cells (pDC) only comprise a minute fraction of human mononuclear leukocytes, but are important anti-viral responders that mediate both innate as well as adaptive immune responses. Persistent activation of pDC enhances HIV pathogenesis by promoting immune suppressive mechanisms such as T regulatory cells. It is therefore important to identify the sources of pDC activation in the context of HIV infection. HIV-associated disruption of gut mucosa associated lymphoid tissue introduces normal flora-lipopolysaccharide (LPS) into systemic circulation, which exacerbates HIV-induced immune activation. We report here that pDC are capable of mediating functional TLR4 signaling upon LPS stimulation, and that pDC of HIV-infected individuals have enhanced TLR4 expression compared to healthy individuals. How TLR4 signaling affects pDC function in HIV infection has not been examined before. Hence we examined the influence of TLR4 signaling in presence of HIVstimulation on pDC and found that it not only potentiated HIV-induced activation but also strongly up-regulated Programmed death ligand-1 (PD-L1) expression and Interleukin-6 synthesis. TLR4 signaling specifically up-regulated PD-L1 expression on activated pDC in presence of HIV stimulation. LPS and HIV co-stimulated pDC demonstrated enhanced migratory potential and repressed T cell proliferation. Together, these results suggest that in the setting of HIV infection enhancement of pDC immune suppressive mechanisms such as PD-L1 may be an outcome of HIV-associated immune activation potentiated by TLR4 signaling.

## Introduction

Cells of the innate immune system express variety of cytoplasmic and membrane-bound pathogen recognition receptors (PRRs) such as, C-type lectin receptor, Retinoic acid-inducible gene1-like receptor, Nucleotide-binding oligomerization domain receptor and Toll-like receptor (TLR). PRRs enable cells to recognize pathogen-associated molecular patterns and mount appropriate immune response against invading pathogens. Among all TLR pathways, the TLR4 signaling pathway is best characterized. TLR4 recognition of its ligand, LPS, activates signaling cascades that mediate cellular defense functions. Depending upon requirement of the adaptor molecule, myeloid differentiation primary response protein 88 (MyD88), TLR4 signaling pathway has two flavours [1–3]. MyD88-dependent pathway operates from cellular surface and results in synthesis of proinflammatory cytokines, while the MyD88-independent pathway signals exclusively from cellular endosomes and results in synthesis of type I IFNs [4]. The MyD88-independent signaling pathway, is mediated with aid of CD14 [5].

Enhanced expression and responsiveness of TLRs by antigen presenting cells has been reported to contribute to hypersystemic immune activation (IA) associated with HIV infection [6–13]. In HIV infected individuals, HIV-associated IA is manifested by elevated levels of immune activation markers on T cells, B cells, monocytes and NK cells, as well as enhanced levels of pro-inflammatory cytokines such as tumor necrosis factor alpha (TNF-α), Interleukin-6 (IL-6) and Interleukin 1-β (IL-β) [9, 13–20]. Although comprising only 0.2-0.5 % of the total human PBMC, pDC help shape the innate as well as adaptive immune responses and are potent anti-viral mediators [21–24]. Because pDCs express CD4, CXCR4 and CCR5 they are susceptible to HIV infection [25]. Researchers have identified that pDC are depleted in HIV infection [26–33], additionally, remnant pDC demonstrate altered capacity to induce T cell proliferation [34] and /or demonstrate poor responsiveness to TLR 7/9 stimulation despite anti-retroviral therapy [35, 36]. Loss of pDC numbers leads to loss of anti-viral activity mediated by IFN-α and enhances viral replication [37, 38].

As with monocytes and NK cells, HIV infection induces persistently activated phenotype in pDC [36, 39]. Persistent activation of pDC leads to induction of tryptophan-catabolizing enzyme, indoleamine-2, 3-deoxygenase (IDO) which induces T regulatory cells[40, 41]. This attenuates anti-HIV immune responses and limits CD4 T cell proliferation [15, 40, 42, 43]. Chronic activation can also potentially disrupt pDC homeostasis by repetitive recruitment of immature pDC from bone marrow; brining about replicative exhaustion and immunosenescence [23, 44]. Optimal restoration of pDC activation and function is essential for suppression of HIV replication.

Besides IA, another feature of HIV infection that contributes to rapid disease progression is, immune exhaustion (IE). IE in HIV infection is manifested by loss or reduced function of T cells and also loss of T cell regenerative capacity and enhanced expression of molecules such as LAG-3, CTLA-4 and programmed death receptor-1 (PD-1) [45–50]. Enhanced PD-L1 expression has been reported on peripheral blood pDCs and mDCs of HIV-infected patients compared to uninfected individuals [51]. PD-1 engagement to its ligand, PD-L1, mediates signaling that suppresses anti-viral as well as anti-tumor CD8^+^ T cell responses [32, 52–60]. The influence of TLR4 signaling on the same has not been examined.

PDC are popularly known for their relatively high expression of TLRs, 7 and 9, while their expression of TLR4 is controversial [61–64]. Our lab has shown that highly purified pDC express TLR4 and that LPS rapidly enhances pDC TLR4 expression [65]. In this study, we reassert these findings and also demonstrate CD14 expression in pDC. We also demonstrate that pDC are capable of inducing phosphorylation and translocation of NF-ΚB to nucleus upon LPS stimulation. This proves that pDC are capable of mediating functional TLR4 signaling in pDC.

HIV-associated IA persists despite Antiretroviral therapy-mediated viral suppression [17, 66–68], indicating that virus independent factors may be involved in its persistence. Microbial products in systemic circulation of HIV-infected individuals have been recently recognized as major mediators of HIV-associated immune activation [15, 18, 69–79]. In their landmark 2006 study, Brenchley and colleagues identified positive correlation between circulating plasma LPS and enhanced innate and adaptive immune system activation [69, 80]. We have observed that viremic HIV-infected individuals have elevated TLR4 expression compared to healthy controls.

Since LPS contributes to HIV-associated IA and HIV-associated IA negatively impacts pDC function, in this study we examined the effect of LPS-induced TLR4 signaling on pDC function in presence of HIV stimulation. We found that TLR4 signaling potently enhanced HIV-induced activation as well as PD-L1 expression on pDC. TLR4 signaling in presence of HIV stimulation simultaneously induced activation and PD-L1 expression on pDC. We evaluated the impact of LPS-modulation on pDC migratory potential and capacity to induce T cell proliferation and found that in presence of HIV stimulation, TLR4 signaling enhanced their migratory potential but repressed their capacity to induce T cell proliferation. Our study identifies an as yet unexamined role ofTLR4 signaling that can enhance pDC contribution to HIV infection.

## Material and Methods

### PBMC preparation and cell enrichments

Institutional Review Board of the New Jersey Medical School approved this study. PBMC were isolated from peripheral blood of healthy or HIV-infected individuals by Ficoll density-gradient centrifugation and re-suspended to 2 × 10^6^ cells/ml concentration in L-glutamine containing RPMI (*10-040-CV, Corning*) supplemented with 10% heat-inactivated fetal calf serum (Atlanta, GA), 25mM HEPES (*H3537-1L, Sigma*) and antibiotics.

Enriched pDC populations, were isolated from PBMC using pDC negative selection enrichment kits *(kit I – 130-092-207 or kit II – 130-097-415 Miltenyi Biotech)*. PBMC resuspended in MACS buffer (0.5 % BSA and 2mM EDTA) were labeled with non-pDC-biotin and microbead antibody cocktail and passed through MACS LD column. Non-pDC cells were retained in magnetic column while “untouched”, purified pDC (>95%) flowed through. CD4 T cells and monocytes were positively selected from PBMC utilizing CD4 and CD14 microbeads respectively (*130-045-101, 130-050-201, Miltenyi Biotech,* respectively).

PBMC or enriched pDC were stimulated with the following activators at the stated concentrations: Ultra-pure LPS (*tlrl-peklps, Invivogen*) or ultra-pure smooth LPS (*tlrl-3pelps, Invivogen*) at 200ng/ml, HIV-1_MN_ and AT-2 inactivated HIV-1_MN_ (Dr. Jeff Lifson, SAIC, NIAID) at 500ng of p24 equivalents, and HSV-1 strain 2931 MOI of 1. TLR4 antagonist, CLI-095 *tlrl-*(*tlrl-cli95, Invivogen*) dissolved in dimethyl-sulfoxide (DMSO)1µg/ml.

### Immunofluorescent staining and flow cytometric analysis

2 × 10^6^ cells/ml PBMC were stimulated for appropriate incubation periods, subsequently washed with 0.1% BSA-PBS and labeled with fluorescently-conjugated monoclonal antibodies for anti human-CD123, BDCA2, HLA-DR, CD14, CD40, CCR7 (*BioLegend*), PD-L1 (*BD Pharmingen*) and TLR4 (eBioscience). Surface-stained cells were then fixed with 1% paraformaldehyde (PFA) and stored at 4°C overnight. For intracellular staining cells were either permeabilized with 0.5% Saponin or 0.1% Triton and stained for TNF-α and/or IL-6 (*BioLegend*). Cells were washed with 2% serum supplemented 1X PBS and subsequently fixed with 1% PFA. 300,000 cellular events were recorded on BD LSR II for non-enriched cultures, and pDC were identified based on co-expression of CD123 and BDCA-2. For enriched pDC cultures, 10,000 events were recorded. Data collected as FCS files were then analyzed with FlowJo ® 9.2 software (Treestar Inc.).

Phosphorylation of NF-ΚB (p65) was evaluated utilizing BD Phosflow assay. Briefly, cells were stimulated and then surface-stained for pDC markers: CD123 and HLA-DR on ice. Because BDCA2 cannot withstand ethanol permeabilization, it was excluded as pDC marker in this assay. Cells were subsequently washed and fixed with 2% PFA on ice for 30 minutes, followed by wash with 1X PBS and subsequently permeabilized with 70% ethanol overnight at −20 °C. Cells were then intracellularly-stained for NF-ΚB (p65) (*BD Biosciences)* and acquired on the BD LSR II.

*Measurement of NF-*Κ*B translocation with Amnis ImageStream*

4 × 10^6^/ml PBMC were stimulated with LPS and subsequently stained for BDCA2, CD123 and CD14. Following permeabilization with 0.1% Triton, cells were intracellularly-stained with anti-NF-ΚB (p50) *(BioLegend).* Nuclear stain DRAQ5 (*Molecular Probes*) was added right before samples were acquired on Amnis ImageStreamX Mark II multispectral flow cytometer. 1000-3000 pDC events and analyzed utilizing IDEAS® software (*Amnis*).

### Quantitative RT-PCR

Gene expression of TLR4 and CD14, was evaluated by quantitative real-time PCR. Total 200,000 enriched pDC and monocytes from same donor were incubated with ultra-pure LPS. Total RNA was subsequently harvested from RLT-lysed cells utilizing RNeasy® Plus micro kit (*74034, Qiagen*). The cDNA copy of RNA (4 µl) was reverse-transcribed using random hexamers. Concentration of RNA and cDNA was measured using the NanoDrop 1000 spectrophotometer (*ND 1000 V3.3.0, Thermo Fischer Scientific*). Real-Time PCR was performed on an Applied Biosystems 7500 Real Time PCR system. DNA templates were amplified utilizing highly specific primer probes (Taqman gene expression assay (20X)) for CD14 (Hs02621496_s1), TLR4 (Hs00152939_m1), and β-actin (Hs99999903_m1) and TaqMan gene expression Master Mix (2X) (*1310185, Life Technologies*). PCR cycle conditions utilized were as follows: 50°C 2 minutes, 95°C 10 minutes, 95°C 15 seconds, 62°C 1 minute. Expression of target genes relative to housekeeping gene control and calibrator (baseline non-treated sample, normalized to housekeeping gene) were calculated utilizing 2^ΔΔ-Ct^ formula [81].

### Transwell migration assay

Total 1×10^5^ enriched pDC / sample, were stimulated with ultra-pure LPS in the presence or absence of HIV_MN_-AT-2. PDC were then washed and placed in the upper chamber of 5 µm pore size Transwells (3421, *Corning*) for 2h at 37°C. Some bottom chambers of the Transwell contained media supplemented with 100ng/ml recombinant human CCL19/MIP-3β (*361-MI-025, R & D systems*). Media without chemokine served as negative control. Subsequenlty, pDCs from individual bottom chambers were rinsed and stained for pDC markers and placed into TruCount beads tubes (*340334, BD Biosciences*) to calculate absolute number of migratory pDC using formula ((# of events in region containing cells)/(# of events in region containing beads)) × ((# of beads per test)/(test volume)).

### CFSE-dilution assay

Allogeneic CD4 T cells were isolated from PBMC using a positive-selection T cell isolation kit and labeled with 5mM/ml carboxyfluorescein diacetate succinimidyl ester (CFSE) as per protocol (*C34554, Molecular Probes, Life Technologies*). Enriched pDC were exposed to LPS with or without HIV_MN_ AT-2 stimulation in the presence of 10ng/ml Interleukin-3 (IL-3) for fifteen hours. Post-stimulation pDC were co-incubated with CFSE-labeled T cells at a 1:20 stimulator:responder ratio (i.e., 5000 pDC:100,000 T cells) for 6 days. T cells stimulated with 200 IU/ml Interleukin-2 (IL-2) served as negative control, while T cells stimulated with 9µg/ml phytohemagglutinin (PHA) (*HA15, Remel Europe*) and IL-2 served as positive control for T cell proliferation. T cells were then harvested and stained for CD3 and CD4 and subsequently assessed for CFSE dilution.

### Statistical analysis

Data from repeat experiments were examined for statistical significance with repeated measures, one-way ANOVA with Bonferroni post-hoc test and selected pair comparisons using GraphPad Prism software. In some instances if data was not normally distributed,

Friedman test was used. Student’s paired, unpaired or non-parametric t-test was utilized wherever appropriate. In some instances data was normalized or transformed by deriving the logarithmic value of data. If statistics determined this way was applied back to original, non-transformed data then the data was represented with geometric means and 95% confidence intervals. * p<0.05, ** p<0.01, *** p<0.001.

## Results

### Gating strategy for identification of pDC from human PBMC

Human pDC are high co-expressers of IL-3 receptor, CD123, and C-type lectin, BDCA2 [82]. To identify pDC utilizing multi-spectral flow cytomtery, PBMC were harvested from human peripheral blood, stimulated and subsequently labeled with monoclonal antibodies specific against CD123, BDCA2 and acquired on BDLSR II. Fig. 1A., represents basic gating scheme utilized for identifying pDC population from PBMC. Firstly, PBMC were defined by gating forward scatter and side scatter area (FSC-A and SSC-A, respectively). From gated PBMC, we identified pDC by their high co-expression of CD123 and BDCA2. From the pDC gate, we identified singlet pDC by gating FSC and SSC-width. Specificity of staining antibody was confirmed by utilizing isotype antibodies for respective markers. Typically, pDC constituted 0.2 to 0.5 % of total isolated PBMC. *LPS stimulation enhances TLR4 but down-regulates CD14 expression on pDC* Endosomal TLRs 7 and 9 are expressed by pDC, providing the cell with potent capacity to recognize RNA and DNA viruses [83–86]. Whether pDC express the LPS receptor, TLR4 is controversial, with one study, which utilized poorly purified pDC, claiming that pDC are non-responsive to LPS stimulation and require the aid of mDC [87]. On the other hand, we have demonstrated that pDC express TLR4 at low but inducible levels utilizing highly purified pDC [65]. Other studies have subsequently confirmed our observations [88–90]. In this study we proceeded by confirming TLR4 expression on pDC. We examined TLR4 expression in freshly isolated (baseline) and ultra-pure LPS stimulated pDCs. Ultra-pure LPS is highly purified form of LPS free of TLR2 signaling contaminants and thus only activates TLR4 signaling. In parallel to analyzing TLR4 expression on pDC, we also assessed TLR4 expression on monocytes in this study. PBMC were stained immediately after isolation or after 6h incubation in presence or absence (mock) of LPS stimulation. Immediate staining of pDC and monocytes enabled us to evaluate TLR4 expression in these cells at baseline or constitutive levels. PDC were identified as per the gating strategy explained above. Monocytes were identified by strong CD14 but lack of BDCA2 expression. In agreement with our previous findings, LPS stimulation enhanced TLR4 expression on pDC (Fig. 1B). Monocytes demonstrated higher TLR4 expression compared to pDC in mock state (Fig. 1D). Unlike with pDCs, LPS stimulation did not significantly enhance mock state TLR4 expression on monocytes. Freshly isolated pDC expressed TLR4 although at lower levels compared to monocytes, but LPS significantly enhanced TLR4 expression on pDCs (Fig. 1C and E). Thus, we confirmed our previous findings that pDC express TLR4 and also comparatively assessed the same against monocytes.

**Figure 1.**
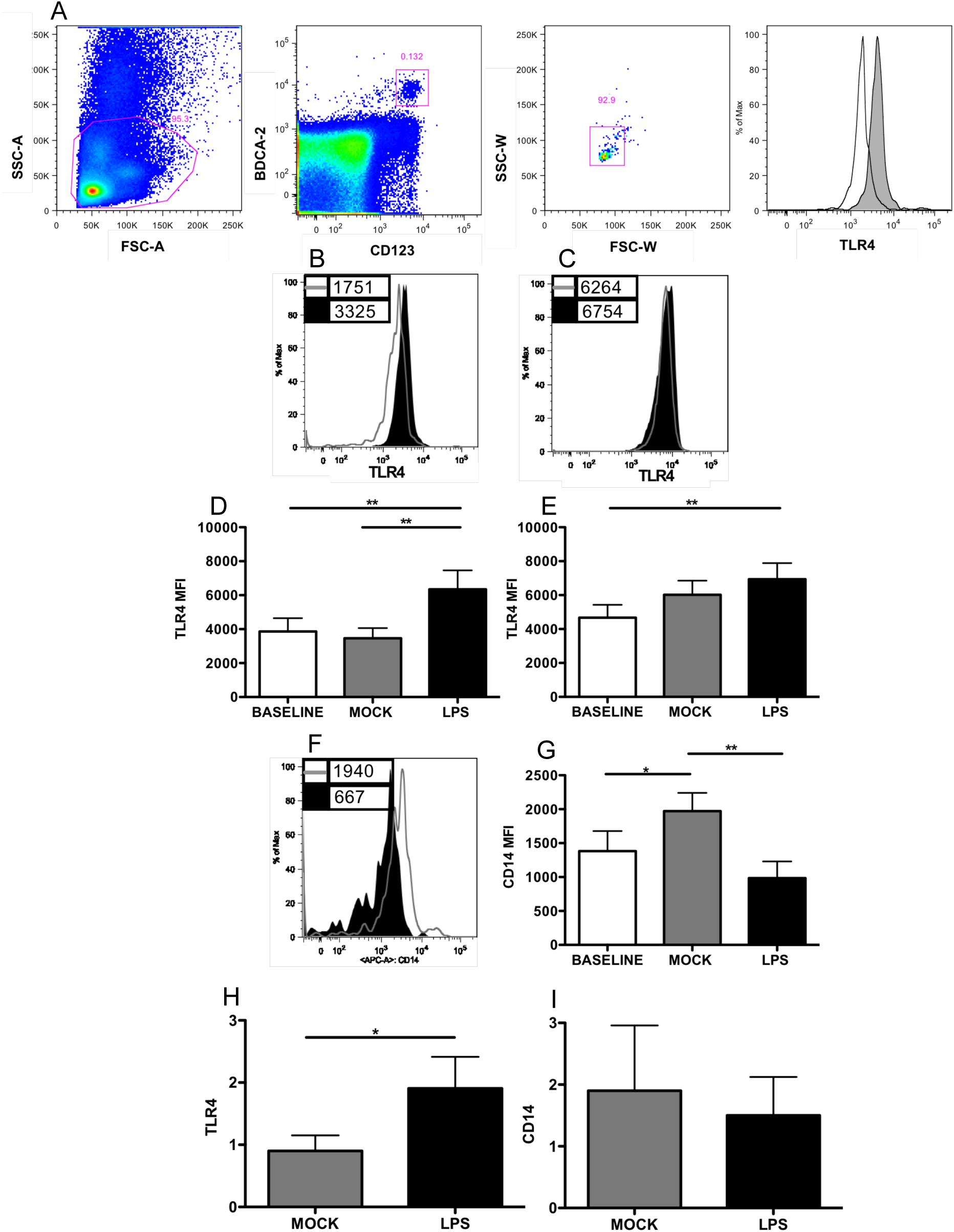
LPS stimulation enhances TLR4 but not CD14 expression on pDC. (A) PBMC were gated by plotting FSC-A vs. SSC-A, subsequently pDC were identified by their co-expression of CD123 and BDCA-2. Singlet pDC (FSC-W and SSC-W) were analyzed for markers of interest. Sample histogram demonstrating singlet pDC expression of TLR4 (filled) vs. isotype (clear) stain. PBMC were either immediately stained with pDC markers, monocyte marker, CD14 and TLR4 or after 6h incubation with or without 200ng/ml ultra-pure LPS. Histogram overlays compare TLR4 expression on pDC (B) and monocytes (D) between 6h mock (gray) and LPS-stimulated sample (black). Legends on histogram represent mean fluorescence intensity (MFI) of TLR4. (C) TLR4 expression is present on freshly-isolated (baseline) pDC and LPS stimulation significantly enhanced baseline as well as mock state TLR4 expression on pDC. (E) LPS stimulation significantly enhanced baseline monocyte TLR4 expression. (F) Histogram overlay represents CD14 MFIs and compares 4h LPS stimulated (black) vs. unstimulated state (gray) CD14 expression on pDC. (G) CD14 expression is present on pDC at baseline and is enhanced in absence of stimulation, but LPS stimulation significantly down-regulates it. (H-I) pDC were purified and stimulated with 200ng/ml ultra-pure LPS or left unstimulated for 2h; message was extracted and mRNA of TLR4 and CD14 was quantified by RT-PCR. LPS induced nearly two-fold enhancement in baseline pDC TLR4 gene expression (H), but CD14 gene expression was unaffected (I). Data represented as mean +/− SEM of TLR4 MFI from three (G-J), five (F) or eight (B-E) independent experiments.

CD14 is a GPI-linked receptor that enables delivery of LPS to MD-2, and enables cellular responses to low levels of LPS [91, 92]. CD14 is capable of TLR4-independent signal transduction involving phospholipase C (PLC)γ2 and subsequently the nuclear factor of activated T cells pathway to initiate DC apoptotic death upon exhaustion and functions exclusively in dendritic cells [93–96]. Whether pDC express the TLR4 co-receptor, CD14 has not been reported. Here we demonstrate that pDC express CD14. We utilized lineage antibody cocktail to exclude conventional high CD14 expressing cells such as monocytes, and examined lineage-negative, singlet pDC. We identified CD14 expression on freshly isolated pDC, and found that it was enhanced in absence of stimulation, but significantly down regulated in presence of LPS stimulation (Fig. 1 F, G).

To evaluate the functional capacity of CD14 expressed by pDC, we examined pDC activation in response to two LPS chemotypes; smooth and rough. Based on absence or presence, of oligosaccharide (O-antigen), LPS can be distinguished as rough or smooth respectively [91]. Smooth LPS can mediate TLR4 signaling only in cells equipped with CD14, while rough LPS can mediate signaling in presence or absence of cellular CD14 [94]. PDC enhanced CD40 expression in response to stimulation with smooth LPS indicate therefore that pDC express CD14 (supplementary fig. 1). In addition, both forms of LPS equally induced pDC activation. These results provide further support that pDC express CD14.

To verify our observations and exclude the possibility of a false–positive result, we corroborated our findings with RT-PCR assay. We evaluated gene expression of both TLR4 and CD14 in highly purified pDC, utlizing highly specific primer probes. We also tested TLR4 and CD14 transcripts in positive-control cells. i.e. monocytes, from the same donor. Monocytes demonstrate elevated TLR4 and CD14 trancript levels compared to pDC (data not shown), but LPS stimulation enhanced baseline TLR4 gene expression in pDC by two fold (Fig. 1 H), while CD14 transcript levels were unaffected (Fig. I). These experiments enabled us to confirm that pDC express TLR4 and helped identify protein as well as transcript level expression of CD14 in pDC.

### TLR4 signaling induces NF-ΚB activation in pDC

The MyD88-dependent TLR4 pathway employs TIR domain-containing adaptor protein (TIRAP) and Interleukin-1 receptor associated kinase 4 (IRAK4) and results in production of pro-inflammatory cytokines through activity of NF-ΚB downstream of TNF receptor-associated factor 6 (TRAF6) [4]. Signal-induced degradation of NF-ΚB inhibitor, IκBα, releases NF-ΚB, which then translocates to cell nucleus and mediates transcription of target genes [97, 98]. We examined the activation of NF-ΚB in pDC upon LPS stimulation by assessing phosphorylation and nuclear translocation of NF-ΚB. After we confirmed that pDC express the LPS receptor, TLR4, we evaluated signaling capacity of the receptor by examining the effect of LPS stimulation on activation of NF-ΚB.

In order to evaluate the phosphorylation of NF-ΚB protein, p65, in response to LPS stimulation we utilized a phospho-flow assay as discussed in methods section. LPS stimulation significantly induced NF-ΚB phosphorylation within twenty minutes in pDC (Fig. 2A). This proved that TLR4 signaling in pDC initiates the first event in NF-ΚB activation. Whether the activated p50:p65 heterodimer of NF-ΚB translocates to the nucleus is absolute measure of its functionality. Hence we examined the translocation of NF-ΚB in pDC following 45 minutes stimulation with LPS using Amnis ImageStream. NF-ΚB translocation in pDC was compared to monocytes, which served as positive control cell type for demonstrating potent TLR4 signaling.

**Figure 2.**
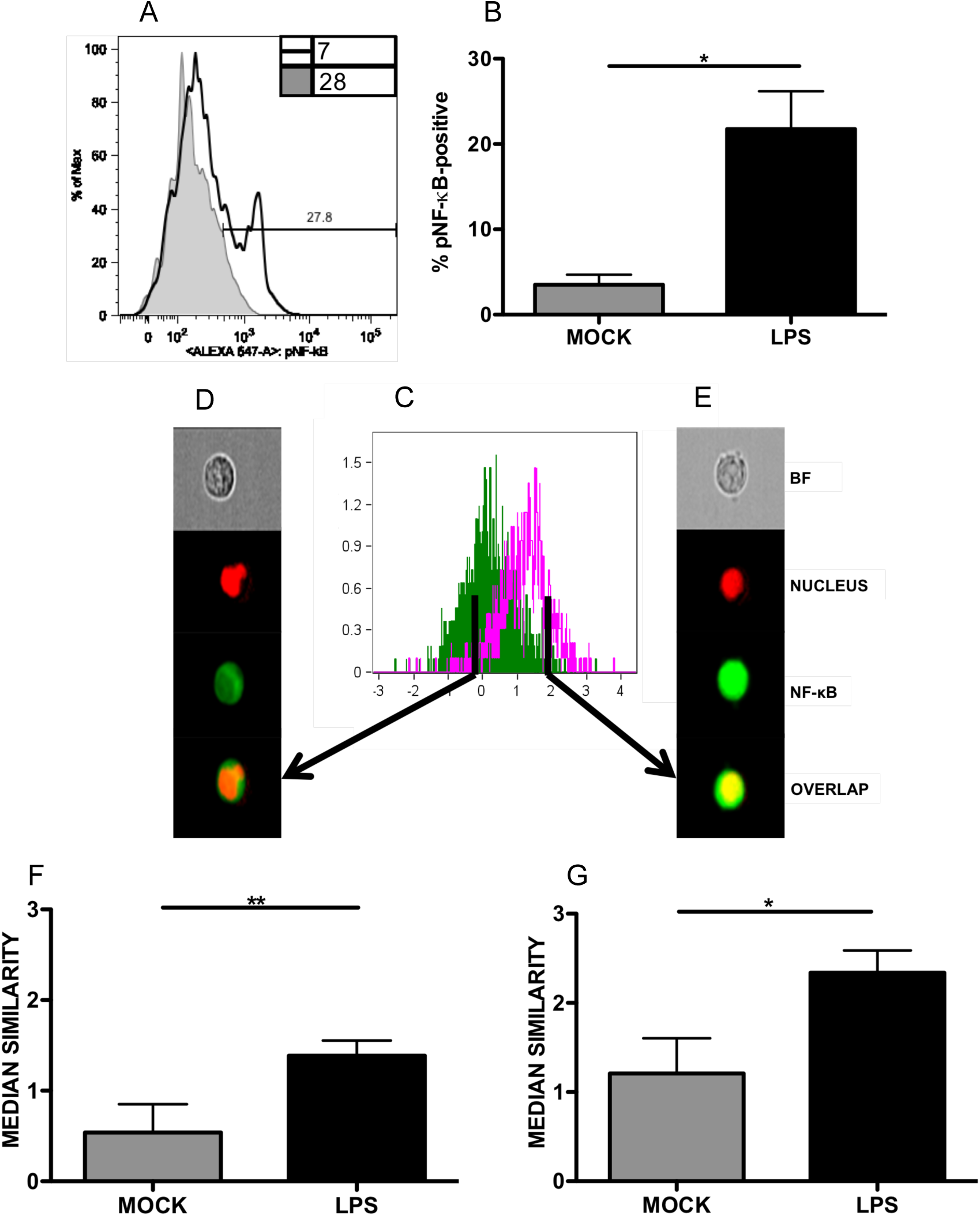
LPS induces rapid phosphorylation and translocation of NF-κB in pDC. Translocation of NF-κB was tested on AMNIS ImageStream. PBMC were stimulated with 200ng/ml ultra-pure LPS for 20 min, before being surface-stained with CD123, HLA-DR and intracellularly-stained for phosphorylated NF-κB (p50). (A) histogram overlay represents LPS-induced (clear) enhancement in frequency of phosphorylated NF-κB-positive pDC compared to mock sample (filled). (C) Overlay of histograms represents NF-κB translocation in pDC upon LPS stimulation (pink) compared to unstimulated state (green). (E) Image plates representing LPS-induced nuclear-translocation of NF-κB in pDC. Translocation is demonstrated by overlap of nuclear stain (red) with NF-κB stain (green) which produced yellow coloring in the nuclear area. Comparison of similarity score medians between unstimulated and LPS-stimulated pDC (F) and monocytes (G) indicated that LPS stimulation significantly enhanced median similarity score in both cell types. Data summary represented as mean of of four (A) or eight (E-F) independent experiments +/− SEM.

Utilizing the gating strategy described in supplementary fig. 2., only sharply focused, uniformly stained pDC and monocytes were examined for translocation of NF-ΚB (p50). Fig. 2 B is a histogram overlay depicting LPS induced nuclear translocation of NF-ΚB (pink) in pDC compared to unstimulated sample (green). Translocated events appeared distinctly yellow, owing to overlap of nuclear (red) and NF-ΚB stain (green), demonstrating nuclear presence of NF-ΚB (Fig. 2C). Similarity score measures overlap of nuclear and NF-ΚB stain. Median of similarity score was utilized to quantify nuclear presence of NF-ΚB. Significant LPS-induced NF-ΚB translocation was not only observed in monocytes but also in pDC (Fig. 2E and F, respectively). Hence, pDC express TLR4 and this receptor mediates functional signaling in these cells.

### TLR4 signaling induces pDC activation and PD-L1 expression

LPS is a commonly utilized stimulus for inducing dendritic cell activation. Activation of pDC is manifested by up-regulated expression of CD40. HIV-associated immune exhaustion is manifested on antigen presenting cells by up-regulated PD-L1 expression. During HIV infection, PD-L1-expressing APCs repress cytotoxic anti-HIV responses mediated by CD8 T cells. We therefore examined the influence of LPS on pDC expression of PD-L1 to evaluate whether LPS could potentiate pDC capacity to induce immune exhaustion. We have observed that LPS stimulation enhances pDC expression of CD40 and PD-L1. To ascertain that this effect of LPS stimulation is indeed mediated through TLR4 signaling, we utilized the TLR4 antagonist, CLI-095. The antagonist inhibited signals mediated through the intracellular domain of TLR4 receptor, without disrupting LPS binding to TLR4 [99]. PBMC were pre-treated with the antagonist for 3h followed by 6h stimulation with LPS. The antagonist significantly repressed LPS-induced activation and PD-L1 expression by pDC (Fig. 3A-B). These results not only confirmed that pDC respond to LPS by enhancing pDC activation and PD-L1 expression, but also proved that this effect of LPS stimulation is mediated specifically by TLR4 signaling in pDC.

**Figure 3.**
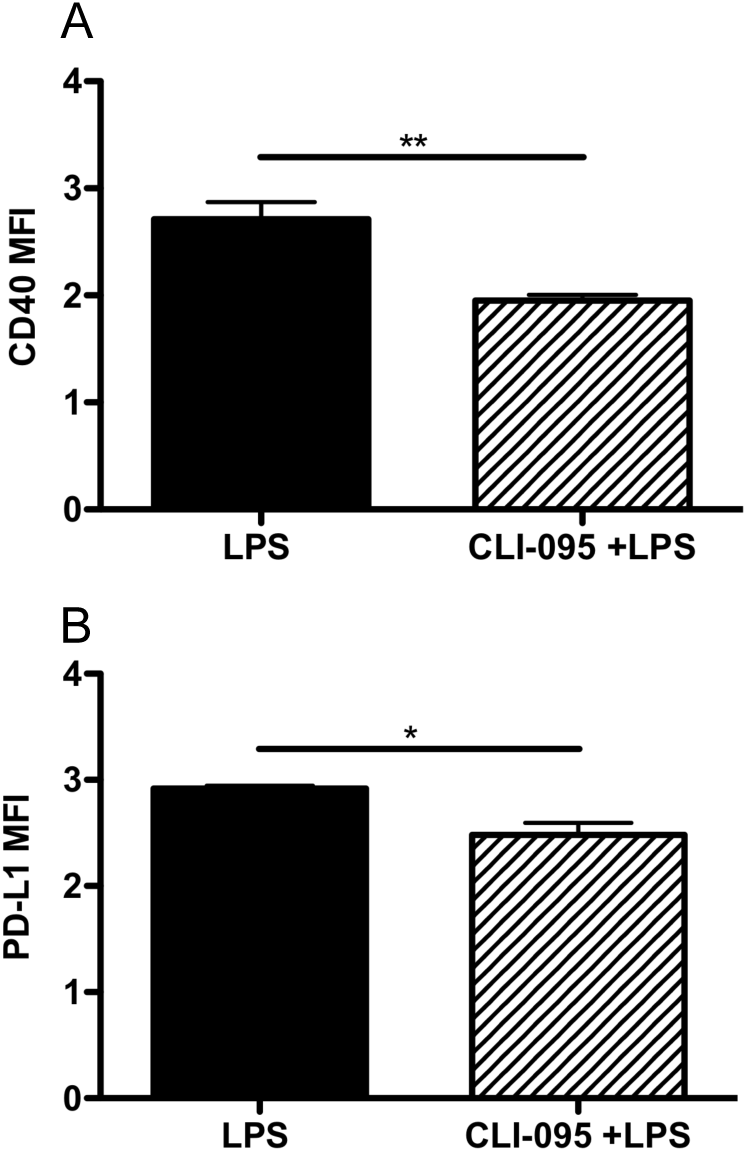
LPS stimulation induces pDC activation and PD-L1 expression through TLR4 signaling. PBMC were pre-treated with 1µg/ml TLR4 antagonist (CLI-095) for 3h prior to stimulation with 200ng/ml ultra-pure LPS for 6h. Cells were subsequently stained with pDC markers, CD40 and PD-L1. LPS-induced pDC expression of CD40 (A) and PD-L1 (B) was reduced by theTLR4 antagonist. Data represented as logarithmic mean +/− SEM of four (for CD40) or three (for PD-L1) independent experiments.

### Viremic HIV-infected individuals demonstrate enhanced TLR4 expression on pDC

HIV infection perpetuates itself through immune activation. For instance, activation of T cells activates the transcription factor NF-ΚB, which encourages transcription of integrated virus [97, 100]. Enhanced expression and responsiveness of TLRs by APCs has also been reported [6–8]. If pDC up-regulate TLR4 expression in setting of HIV infection, then it may affect its responsiveness to LPS induced immune activation. We therefore examined if there were differences in TLR4 expression between pDC of healthy and HIV-infected individuals. Freshly isolated pDC from healthy and HIV-infected individuals were stained for pDC markers and TLR4. When we compared baseline TLR4 expression on pDC of HIV-infected individuals unstratified for viral load (n=55) and healthy controls we found distinct trend for higher TLR4 expression in the former versus the latter (data not shown). Since most of our patient populations are virally suppressed, we examined TLR4 expression solely in viremic HIV-infected individuals (viral load >400 copies/ml) (n=17), to exclude possibility of viral suppression biasing our observation. HIV-infected individuals with high viral load demonstrated significantly elevated TLR4 expression compared to healthy controls (HCs) (Fig. 4A). High viremia may influence TLR4 expression on pDC in HIV infection. Monocytes of HIV-infected individuals also demonstrated distinct trend for up regulated TLR4 expression compared to HCs (data not shown).

**Figure 4.**
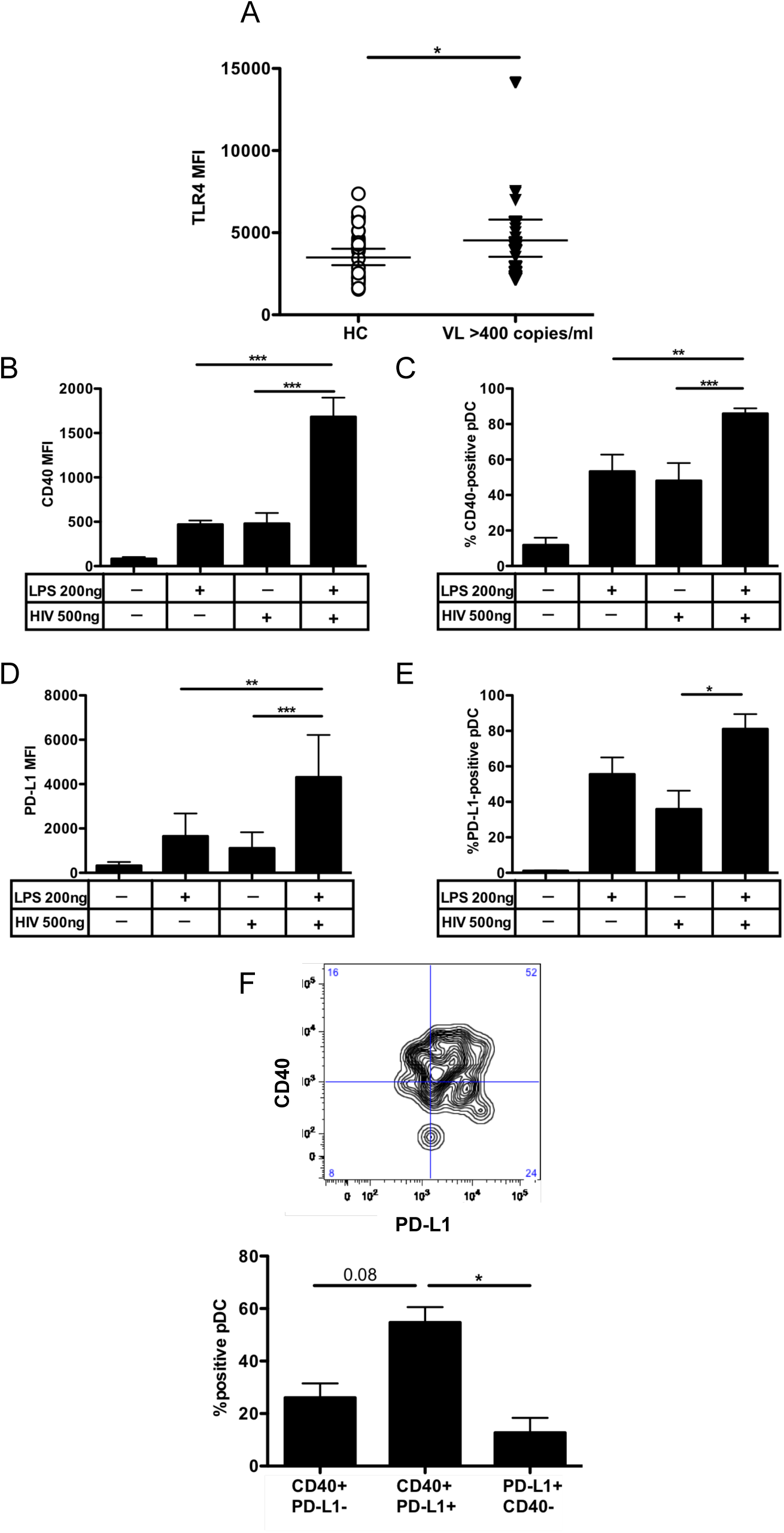
LPS enhances HIV-induced pDC activation and PD-L1 expression. Freshly-isolated PBMC from healthy controls and viremic HIV-infected individuals (viral copies (VL) >400 copies/ml) were stained with pDC markers and TLR4. (A) pDC from viremic HIV-infected individuals demonstrated higher TLR4 expression compared to healthy controls. (B-E) PBMC were pre-treated with 200 ng/ml ultra-pure LPS for 2h prior to stimulation with HIV_MN_ for 8h. Cells were subsequently stained with pDC markers, CD40 and PD-L1. LPS enhanced HIV-induced CD40 expression (B) and percentage of CD40-positive pDC (C). LPS also enhanced HIV induced PD-L1 expression on pDC (D). LPS and HIV in combination also induced more PD-L1-positive pDC than virus treatment alone (E). (F) Sample dot plot compares CD40 vs. PD-L1. TLR4 signaling in presence of HIV stimulation significantly enhanced frequency of PD-L1,CD40 double-positive pDC. Data represented as mean +/− SEM (B,C,E and F) or geometric mean with 95% confidence interval (A and D) of three or eight independent experiments.

### TLR4 signaling in pDC enhances HIV-induced activation and PD-L1 expression

HIV infection is associated with persistent immune activation and immune exhaustion. How innate cells such as pDC are affected and/or contribute to pathogenesis in this setting is not clear. We observed that TLR4 signaling in pDC enhanced expression of both CD40 and PD-L1. In order to examine how TLR4 signaling in presence of HIV stimulation may influence pDC activation and PD-L1 expression, we co-stimulated pDC with LPS and HIV. We primed pDC with LPS for 2h followed by HIV_MN_ stimulation for 8h. Our findings indicate that neither treatment by itself could induce pDC activation and PD-L1 expression to the extent as LPS and HIV in combination (Fig. 4B and D). The combination treatment also enhanced frequency of activated and PD-L1-positive pDC (fig. 4C and E). This indicates that LPS-mediated TLR4 signaling can potently enhance HIV-induced pDC activation and PD-L1 expression. This may be an outcome of additive signaling from TLRs 4 and 7. We also examined the effect of TLR4 signaling on expression of positive costimulatory molecules, CD80 and CD86, but did not observe TLR4-induced potentiation of HIV-induced upregulation of these molecules (data not shown).

We next examined if LPS and HIV in combination might induce PD-L1 expression simultaneously or separately on activated pDC. For this purpose, pDC were primed with LPS for 2h followed by HIV_MN_ stimulation for 6h. Subsequently, the cells were stained for pDC markers, CD40 and PD-L1. Utilizing FlowJo software a contour plot comparing PD-L1 versus CD40 expression was generated from the pDC population for all samples. We compared frequency of single or double-positive pDC, from each quadrant of the dot plot and found that the percentage of CD40, PD-L1 double positive pDC was significantly enhanced with LPS and HIV combination treatment compared to individual HIV or LPS treatment (Fig. 4F). LPS treatment induced PD-L1-single-positive pDC, but in combination with virus, LPS induced PD-L1 expression specifically on activated pDC. In HIV infection, as the components of gut microflora translocate to systemic circulation, they contribute to HIV-associated immune activation [3,7,27–29], mainly by enhancing levels of cytokines such as pro-inflammatory cytokines IL-6, TNF-α and IL-β [69]. These cytokines participate in refueling the cascade of immune activation [13]. It has been reported that monocytes of HIV-infected individuals produce high levels of pro-inflammatory cytokines due to over-activation induced by systemic LPS [6, 8]. TLR9 stimulation in pDC can induce synthesis of pro-inflammatory cytokines by pDC [101] but whether LPS can can do the same has not been explored. We treated PBMC with LPS for 6h and examined IL-6, TNF-α-positive pDC by intracellular flow cytometry. In parallel to this we examined HSV-induced IL-6, TNF-α-positive pDC. LPS induced TLR4 signaling significantly enhanced frequency of IL-6-positive pDC. LPS demonstrated potency equal to HSV for this (supplementary fig. 3A). TLR4 signaling unlike TLR9 signaling did not in enhance frequency of TNF-α-positive pDC (supplementary fig. 3B). We concluded that LPS stimulation, besides enhancing pDC activation and PD-L1 expression, triggers their selective production of pro-inflammatory cytokine, IL-6.

IL-6 is a known contributor to HIV disease progression and is strongly correlated with increased risk of mortality, non-HIV-related morbidity and immune senescence [13, 102, 103]. By encouraging T cell activation, IL-6 aids in generation of new targets for viral replication [13]. To examine the effect of TLR4 signaling on IL-6 synthesis in presence of HIV stimulation, we primed pDC with LPS for 2h followed by stimulation for 6h with HIV_MN_. Cells were then stained for pDC markers and after permeabilization stained for IL-6. Stimulation with LPS and HIV in combination significantly enhanced pDC frequency of IL-6-positive pDC (Fig. 5A-B). These results support a potential role of LPS-mediated TLR4 signaling in enhancing pathogenesis of HIV infection by enhancing synthesis of IL-6.

**Figure 5.**
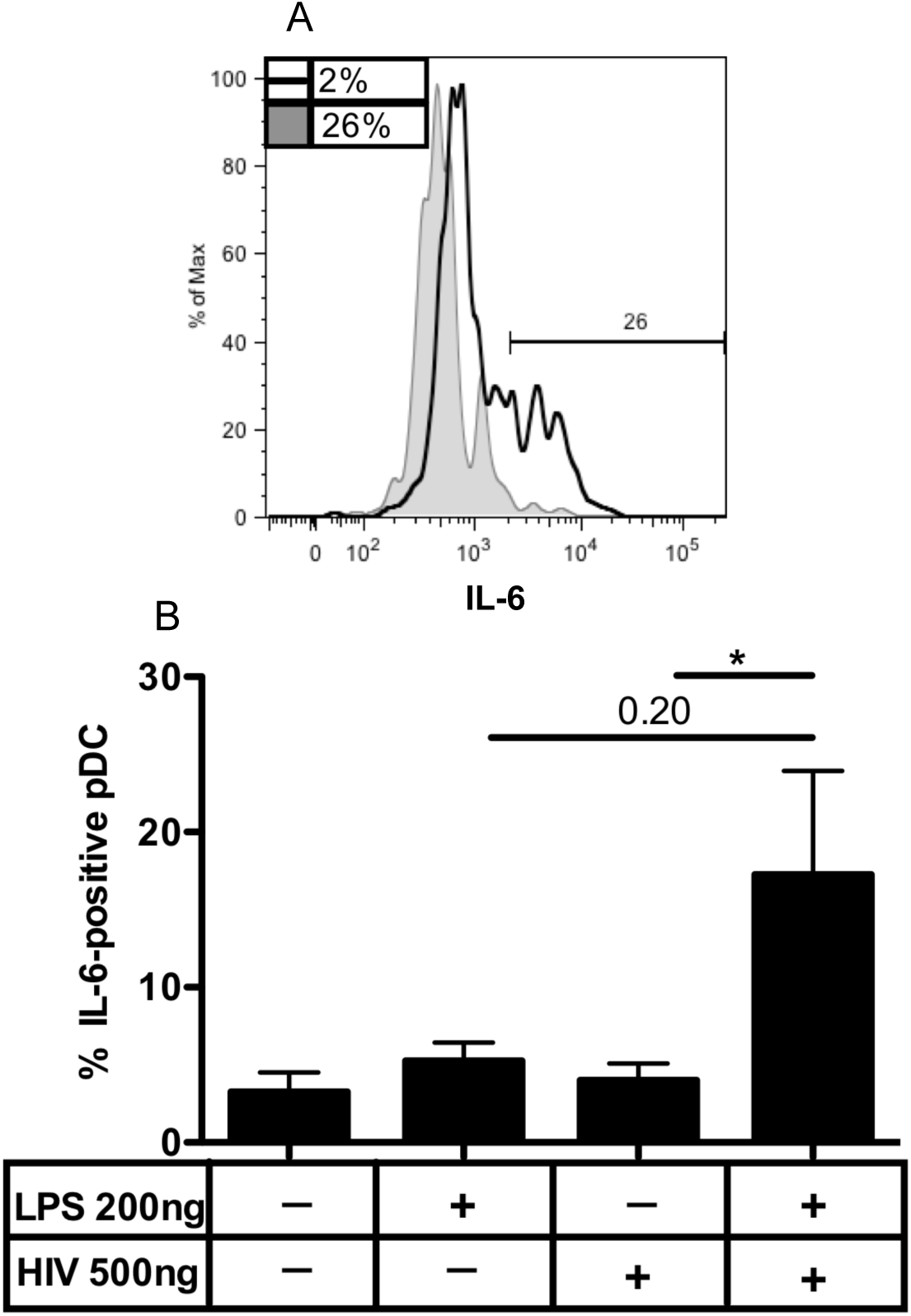
TLR4 signaling in presence of HIV stimulation enhances Interleukin-6 production by pDC. PBMC were pre-treated with 200ng/ml ultra-pure LPS for 2h prior to stimulation with HIV-MN for 6h. Cells were subsequently stained with pDC markers and intracellularly stained for IL-6 after cell permeabilization. (A) Histogram compares frequency of IL-6-positive pDC induced by mock (filled) and combination treated (clear) pDC. (B) TLR4 signaling in presence of HIV stimulation enhanced IL-6 synthesis by pDC. Data represented as mean +/− SEM of four independent experiments.

### LPS and HIV co-stimulated pDC suppress T cell proliferation

TLR4 signaling in pDC in presence of HIV stimulation resulted in enhanced CD40 and PD-L1 expression. To assess the influence of pDC modulated by LPS and HIV treatment on T cell proliferation, we utilized CFSE dilution assay. To ensure that only pDC influence on T cell proliferation was assessed, we depleted non-pDC from PBMC by pDC enrichment. PDC were then stimulated with LPS or HIV_MN_ individually or in combination for 15h and then placed in co-culture with CFSE-labeled allogeneic CD4 T cells. T cells treated with PHA were utilized as positive control for T cell proliferation, while T cells treated with IL-2 served as negative control for T cell proliferation. The co-cultures were allowed to process for 6 days and cells were subsequently harvested, stained for T cell markers (CD3 and CD4) and examined for CFSE dilution by flow analysis. T cells, when co-cultured with HIV and LPS-stimulated pDC, demonstrated repressed T cell proliferation compared to pDC stimulated only with LPS (Fig. 6B-C). We concluded that LPS and HIV co-stimulated pDC exhibit a functionally suppressive phenotype that dampens T cell proliferation, consistent with their expression of PD-L1.

**Figure 6.**
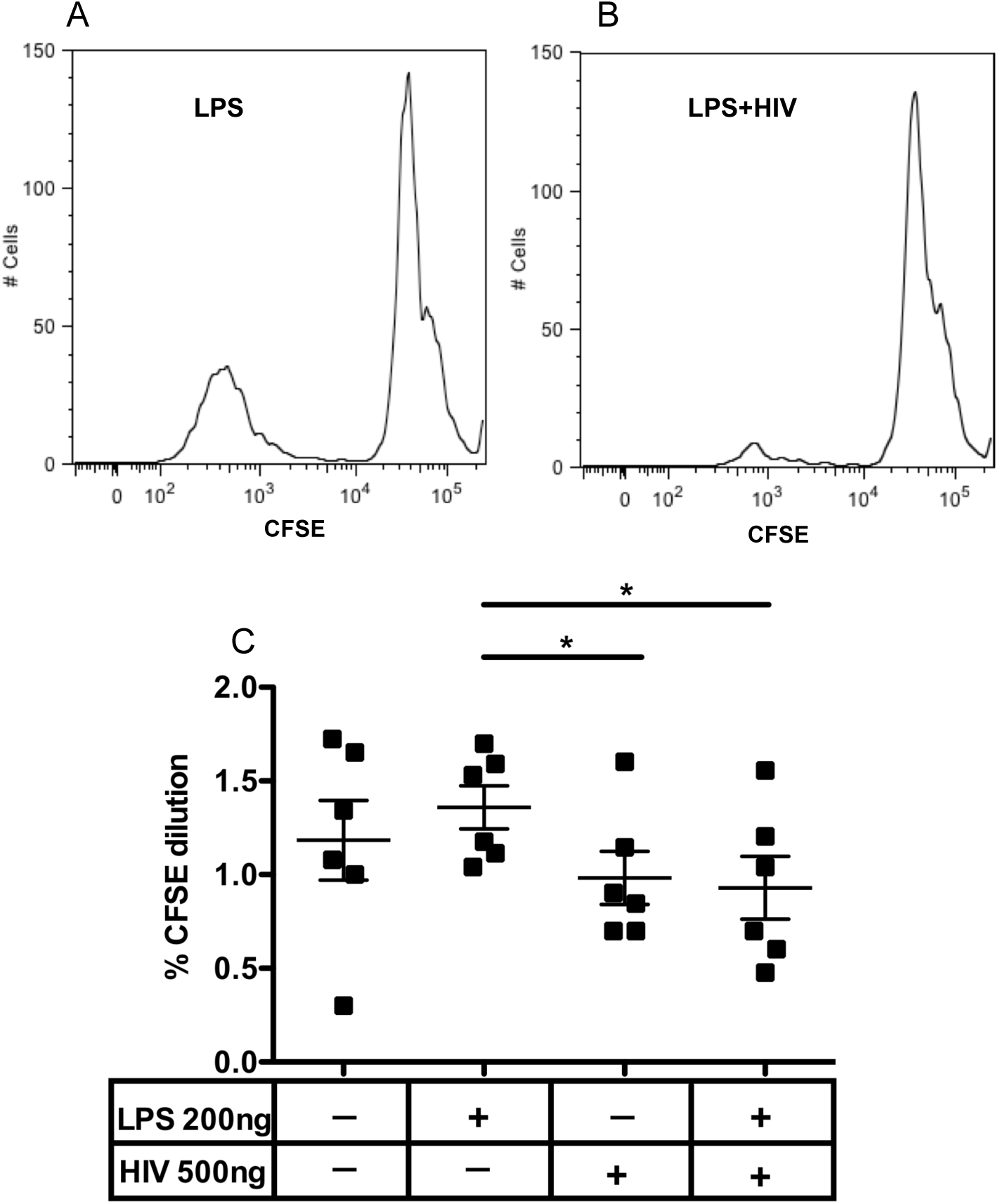
HIV represses T cell proliferation in presence of TLR4 signaling. Enriched pDC were stimulated with LPS, AT2-inactivated-HIV_MN_ or LPS and HIV in combination in presence of interleukin-3 for 15h before being placed in co-culture with CFSE-labeled allogeneic CD4 T cells. Following six day co-culture, cells were harvested and stained for CD4 and CD3. Proliferation of T cell co-incubated with LPS-stimulated pDC (A) and combination treated pDC (B) is shown. (C) Proliferation of T cells was repressed by LPS and HIV co-stimulated pDC. Data represented as mean +/− SEM of four independent experiments.

### LPS and HIV stimulated pDC enhance migration in response to CCL19

We observed that TLR4 signaling in presence of HIV stimulation enhanced pDC activation. PDC enhance expression of migration molecule, CCR7, upon activation [104]. CCR7 ligand, CCL19, can guide migration of pDC to secondary lymphoid organs. Upon migration to lymph nodes, pDC present antigen and co-stimulatory signals to adaptive immune system components, CD4 and CD8 T cells [105–107]. HIV-1 induces CCR7 expression on pDC in vitro [108] and Herbeuval et al., identified enhanced CCR7 expression on pDC of HIV-positive patients [109]. To examine influence of LPS and HIv combination treatment on pDC migratory potential, we first examined the influence of this combination treatment on pDC expression of CCR7. LPS priming enhanced HIV-induced CCR7 expression (fig. 7D). We also examined the effect of LPS and HIV on pDC expression of CD62L, and observed that it induced shedding of CD62L, which is consistent with enhanced pDC activation (data not shown).

**Figure 7.**
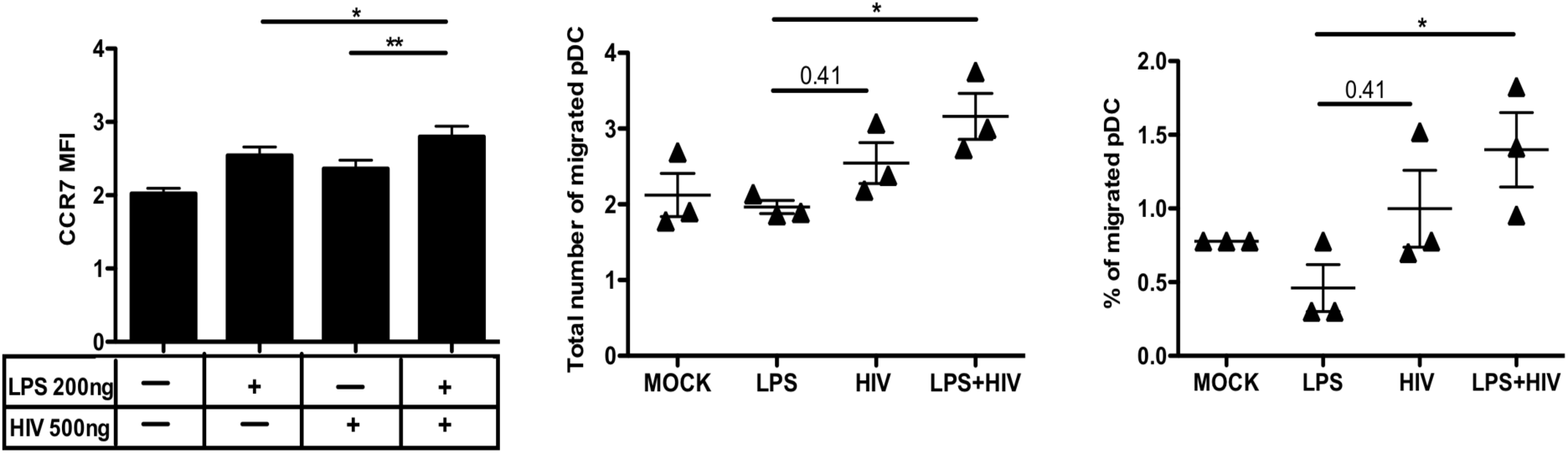
LPS and HIV-stimulated pDC demonstrate enhanced migration in response to CCL19 (A) PBMC were stimulated with LPS, AT2-inactivated-HIV_MN_ or LPS and HIV in combination for 8h. LPS enhanced HIV-induced CCR7 expression on pDC. (B-C) pDC were purified from PBMC and stimulated with LPS, HIV_MN_-AT2 or LPS and HIV_MN_-AT2 in combination for 15 h and then placed in top chamber of Transwell plate for 2h. LPS and HIV co-stimulation enhanced migration of pDC to bottom chamber in response to CCL19. Data represented as mean (logarithm (B-C)) +/− SEM of three independent experiments.

We measured migratory potential of LPS and HIV co-stimulated pDC using Transwell assays. PDC co-stimulated with LPS and HIV_MN_ were placed in top chamber of Transwell plate. Bottom chamber of some wells had CCL19. More pDC co-stimulated with LPS and HIV migrated in response to CCL19 compared to LPS stimulated pDC (Fig. 7B-C). We concluded that LPS and HIV stimulation enhances the frequency of migrating pDC.

## Discussion

HIV infection is characterized by CD4 T cell depletion, uncontrolled viremia, enhanced incidences of opportunistic infections and chronic IA. Persistent IA associated with HIV predicts faster disease progression and is manifested by elevated levels of activation markers on immune cells as well as enhanced expression and responsiveness of TLRs by antigen presenting cells [6–20, 110]. Because TLRs play critical role in facilitating inflammatory responses, their aberrant expression and responsiveness can contribute to HIV-associated IA. In HIV infection TLR activation can be direct, i.e., through virus-mediated TLR7 or 8 signaling or indirect, through TLR signaling mediated by co-infecting pathogens or normal flora products introduced into systemic circulation by microbial translocation [69, 111, 112].

Persistence of IA in HIV-infected individuals is observed in spite of antiretroviral therapy-mediated viral suppression [17, 66–68], indicating involvement of virus independent factors. Gram-negative bacterial LPS is typically identified in plasma of HIV-infected individuals and is known to contribute to HIV-associated IA [68, 69, 73, 74, 113–116]. Besides LPS, other microbial products found in plasma of HIV-infected individuals include bacterial DNA fragments, 16S rRNA [77, 117], and immune system components such as soluble CD14 (sCD14) [69, 113, 115], LPS binding protein (LBP) and anti-LPS antibodies, called endotoxin-core antibodies.

TLR4 signaling can contribute to HIV pathogenesis via multiple modes. For instance, TLR4 signaling has been linked with promoting expression of HIV-1 genome [118]. Plasma LPS can induce persistent TLR4 signaling leading to enhanced pro-inflammatory cytokine production thereby potentiating HIV-associated IA. It has also been demonstrated that HIV glycoprotein induced TLR4 signaling promotes IA in female genital epithelium [119]. In this study, we examined the role of TLR4 signaling in HIV mediated IA, particularly in pDC; a cell type negatively impacted by IA.

TLR4 expression on pDC is controversial, however, we have previously shown that highly purified pDC express TLR4 and that LPS rapidly enhances its expression [65]. Other studies have also confirmed our findings [88–90]. However, the study by Hernandez et al., demonstrated very low MFI for TLR4; even in high TLR4 expressing cells, monocytes. On the other hand we demonstrate high TLR4 MFI (>10^3^) on pDC as well as monocytes even at baseline. The study by Hernandez et al utilized pDC enriched by positive selection. This method of pDC purification induces BDCA2 cross-linking which we have reported affects pDC function [120]. In this study we specifically utilized negatively selected, “untouched” pDC to demonstrate TLR4 transcript levels in pDC. Additionally, we utilized highly specific primer probes to quantify TLR4 gene expression in freshly isolated, unstimulated and LPS-stimulated pDC.

In this study we also examined if pDC are capable of mediating functional TLR4 signaling upon LPS stimulation. We observed rapid induction of NF-κB phosphorylation in pDC upon LPS stimulation, indicating that it was a direct effect of TLR4 signaling in pDC rather than trans-activation by other cells. However, in the future we can address this more directly by examining NF-κB phosphorylation in highly enriched pDC. In this study we also demonstrated that LPS-induced NF-κB translocation in pDC. We observed TLR4 protein and transcript level expression in pDC and also found that LPS induced both activation and translocation of NF-κB, this proves that pDC not only express TLR4 but also that the receptor is capable of mediating functional signaling in these cells.

In vitro HIV stimulation of pDC has been shown to enhance TLR4 expression and LPS can potentiate IA via multiple modes in HIV infection [88]. Hence TLR4 signaling and HIV may share a coadjuvant relationship that enhances disease progression. We have previously demonstrated that LPS up-regulates IRF-7 expression in pDC in an NF-κB-activation-dependent manner [65]. This may be potential point at which LPS can mediate synergistic effects with virus on pDC activation and function. In this study we demonstrated that freshly isolated pDC of viremic HIV infected individuals have enhanced TLR4 expression compared to healthy individuals, indicating that viremia can influence TLR4 expression on pDC. Although enhanced plasma LPS due to compromised gut barrier in viremic patients can enhance TLR4 expression.

To the best of our knowledge we are the first to show that pDC express TLR4 co-receptor, CD14. To rule out the possibility of artifact due to deposition of soluble CD14 (which is known to be shed from monocytes [121]) on pDC, we confirmed our flow cytometry findings by evaluating CD14 transcript levels in highly-enriched pDC. Additionally, we found that smooth LPS ably induced pDC activation, indirectly indicating that pDC express functional CD14. Also both LPS chemotypes similarly induced pDC activation; indictaing that pDC can mediate TLR4 signaling in presence or absence of CD14.

In the T cell activation process, signal 2, is a crucial point where immune responses can be enhanced or repressed, depending upon the type of signal conveyed. The PD-L1: PD-1 pathway is a part of the B7-CD28 family of pathways that convey signal 2 in the lymphocyte activation process. Under normal circumstances, the PD-L1: PD-1 pathway is important for repressing autoimmunity and maintaining self-tolerance, but because it suppresses T cell responses; viruses, establishing chronic infections manipulate it to blunt anti-viral immunity [122]. In chronic viral infections like HIV-1, CD8 T cells up-regulate PD-1 expression and this characterizes them as “exhausted” [123]. The function of exhausted CD8 T cells in chronic HIV-1 infection is negatively regulated by the PD-1: PD-L1 pathway. HIV-infected individuals demonstrate enhanced PD-L1 expression on their APCs, which correlates directly with viral load, indicating potential effect of the virus on PD-L1 expression [51, 58, 70]. Recently, Planes et al., demonstrated HIV Tat-mediated up-regulation of PD-L1 expression on human mDC, which utilizes TLR-4-dependent mechanisms [124, 125].

Whether LPS can influence the PD-L1 expression on pDC in presence of HIV induced IA has not been investigated before. In our study we demonstrated that TLR4 signaling in presence of HIV stimulation enhanced pDC activation as well as PD-L1 expression. Because PD-L1 was specifically induced on activated pDC by the combination treatment, we concluded that enhanced PD-L1 expression on pDC is an outcome of HIV induced pDC activation potentiated by LPS signaling.

Pro-inflammatory cytokines such as IL-6 and TNF-α are elevated in HIV-infected individuals. IL-6 contributes to HIV disease progression and is strongly correlated with increased risk of mortality, immunosenescene and non-HIV-related morbidity. We found that although LPS and virus individually induced IL-6 synthesis by pDC, the individual treatments were not as potent as virus and LPS in combination. This indicates that LPS and HIV may mediate this effect collaboratively.

It is already established that chronic activation of pDC contributes to HIV pathogenesis by triggering apoptosis of CD4 T cells and disrupting pDC homeostasis or by inducing immune-suppressive mechanisms [126]. Through our study we reveal an as yet unexplored role ofTLR4 signaling in abetting HIV pathogenesis by enhancing immune activation and consequently a counter-regulatory mechanism, PD-Ll.

---

**Supplementary figure 1.**
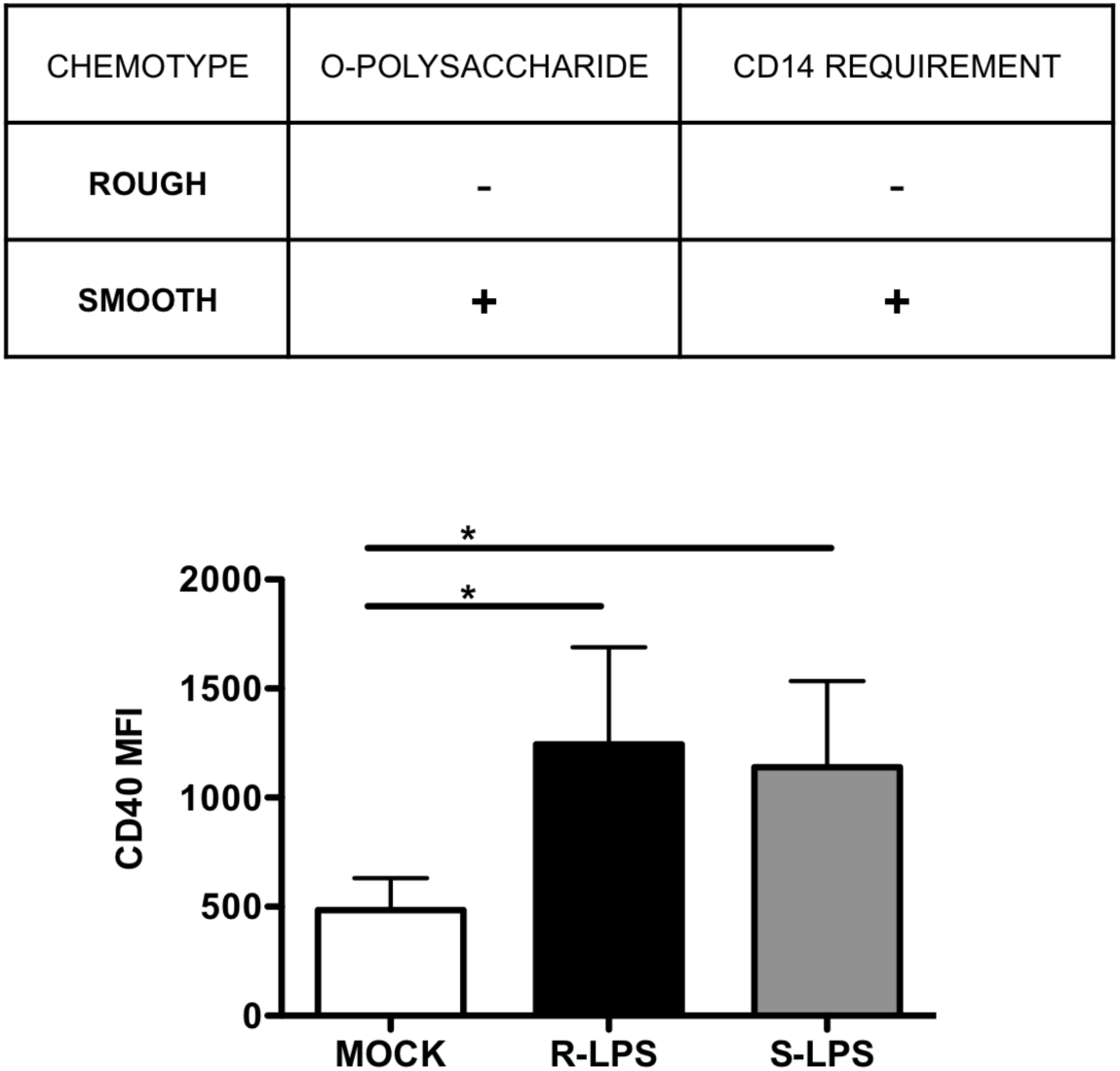
Smooth LPS induces pDC activation. PBMC were either stimulated with 200 ng/ml ultra-pure rough or smooth-LPS or left unstimulated for 6h. Cells were subsequently stained with pDC markers and CD40. Both variations of LPS induced CD40 expression on pDC. Data represented as mean +/− SEM of five independent experiments.

**Supplementary figure 2.**
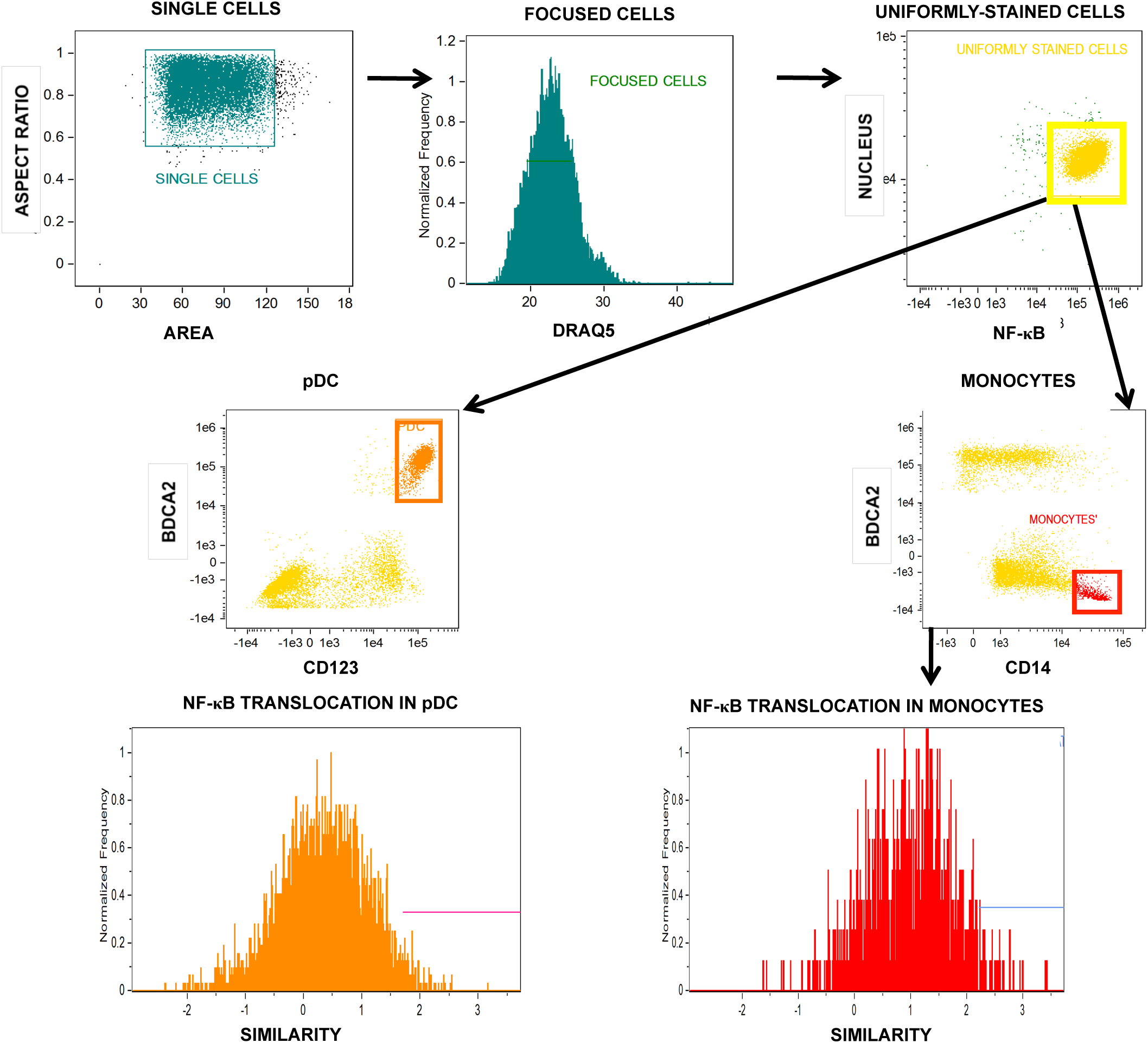
Gating strategy for identifying NF-κB nuclear translocation. Basic gating strategy to examine translocation of NF-κB using AMNIS ImageStream. Single cells were identified by comparing aspect ratio vs. area, sharp images were identified and gated using gradient histogram, cells stained uniformly for nuclear stain (DRAQ5) and NF-κB were gated next. From the uniformly stained cell populations, pDC were identified by their high co-expression of CD123 and BDCA-2. Monocytes were identified by their strong CD14 but lack of BDCA-2 expression. Translocation was computed by Ideas software as correlation between nuclear and NF-κB stain; similarity. Sample histogram of pDC (yellow) and monocyte (red) representing similarity.

**Supplementary figure 3.**
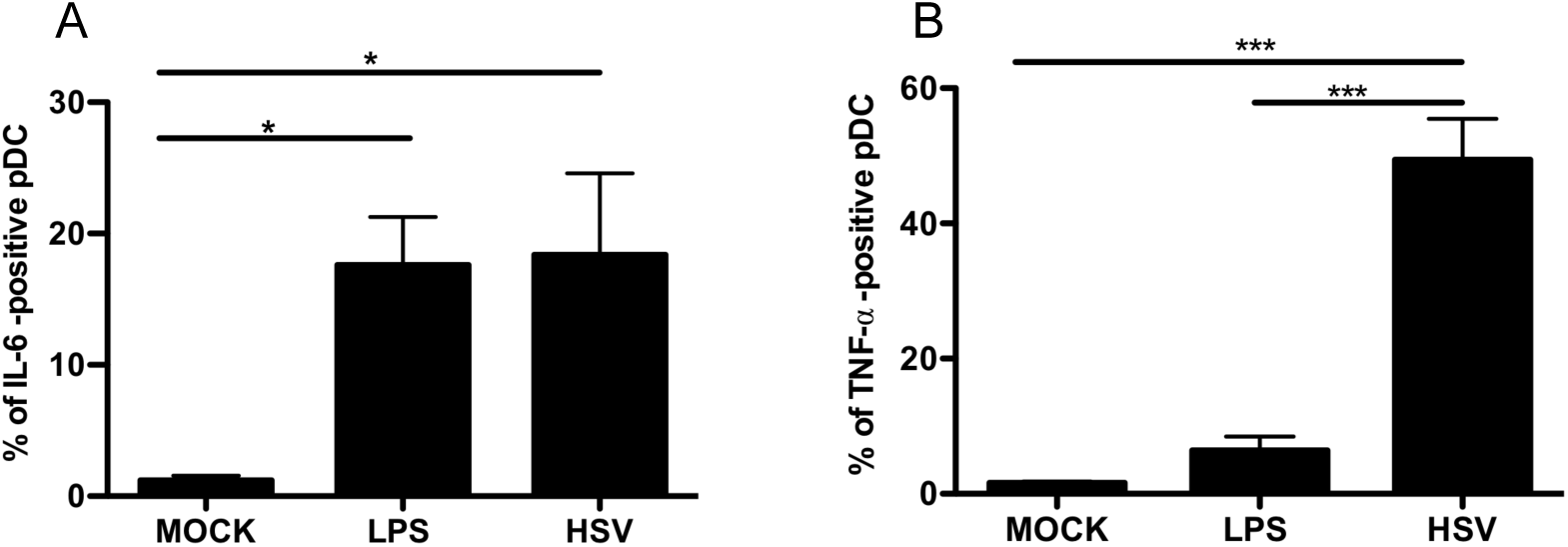
PBMC were pre-treated with 200ng/ml LPS or HSV for 6h. 2h before harvesting cells, Brefeldin A was added. Cells were subsequently stained with pDC markers and intra-cellullarly-stained for IL-6 or TNF-α. (A) LPS and HSV stimulation induced IL-6 synthesis. (B) LPS did not induce TNF-α synthesis. LPS induced far fewer pDC to synthesize TNF-α compared to IL-6. Data represented as +/− SEM of five independent experiments.

## Notes

### Competing Interest Statement

The authors have declared no competing interest.

## References

1. Barton, G.M. and J.C. Kagan, A cell biological view of Toll-like receptor function: regulation through compartmentalization. Nat Rev Immunol, 2009. 9(8): p. 535–42.

2. Kagan, J.C. and R. Medzhitov, Phosphoinositide-mediated adaptor recruitment controls Toll-like receptor signaling. Cell, 2006. 125(5): p. 943–55.

3. Akira, S. and K. Takeda, Toll-like receptor signalling. Nat Rev Immunol, 2004. 4(7): p. 499–511.

4. Hayden, M.S. and S. Ghosh, Shared principles in NF-kappaB signaling. Cell, 2008. 132(3): p. 344–62.

5. Zanoni, I., R. Ostuni, Lorri R. Marek, S. Barresi, R. Barbalat, Gregory M. Barton, F. Granucci, and Jonathan C. Kagan, CD14 Controls the LPS-Induced Endocytosis of Toll-like Receptor 4. Cell, 2011. 147(4): p. 868–880.

6. Lester, R.T., X.D. Yao, T.B. Ball, L.R. McKinnon, W.R. Omange, R. Kaul, C. Wachihi, W. Jaoko, K.L. Rosenthal, and F.A. Plummer, HIV-1 RNA dysregulates the natural TLR response to subclinical endotoxemia in Kenyan female sex-workers. PLoS One, 2009. 4(5): p. e5644.

7. Meier, A., G. Alter, N. Frahm, H. Sidhu, B. Li, A. Bagchi, N. Teigen, H. Streeck, H.J. Stellbrink, J. Hellman, J. van Lunzen, and M. Altfeld, MyD88-Dependent Immune Activation Mediated by Human Immunodeficiency Virus Type 1-Encoded Toll-Like Receptor Ligands. Journal of Virology, 2007. 81(15): p. 8180–8191.

8. Mureith, M.W., J.J. Chang, J.D. Lifson, T. Ndung’u, and M. Altfeld, Exposure to HIV-1-encoded Toll-like receptor 8 ligands enhances monocyte response to microbial encoded Toll-like receptor 2/4 ligands. AIDS, 2010. 24(12): p. 1841–8.

9. Giorgi, J.V., L.E. Hultin, J.A. McKeating, T.D. Johnson, B. Owens, L.P. Jacobson, R. Shih, J. Lewis, D.J. Wiley, J.P. Phair, S.M. Wolinsky, and R. Detels, Shorter survival in advanced human immunodeficiency virus type 1 infection is more closely associated with T lymphocyte activation than with plasma virus burden or virus chemokine coreceptor usage. J Infect Dis, 1999. 179(4): p. 859–70.

10. Giorgi, J.V., Z. Liu, L.E. Hultin, W.G. Cumberland, K. Hennessey, and R. Detels, Elevated levels of CD38+ CD8+ T cells in HIV infection add to the prognostic value of low CD4+ T cell levels: results of 6 years of follow-up. The Los Angeles Center, Multicenter AIDS Cohort Study. J Acquir Immune Defic Syndr, 1993. 6(8): p. 904–12.

11. Liu, Z., W.G. Cumberland, L.E. Hultin, A.H. Kaplan, R. Detels, and J.V. Giorgi, CD8+ T-lymphocyte activation in HIV-1 disease reflects an aspect of pathogenesis distinct from viral burden and immunodeficiency. J Acquir Immune Defic Syndr Hum Retrovirol, 1998. 18(4): p. 332–40.

12. Paiardini, M. and M. Muller-Trutwin, HIV-associated chronic immune activation. Immunol Rev, 2013. 254(1): p. 78–101.

13. Appay, V. and D. Sauce, Immune activation and inflammation in HIV-1 infection: causes and consequences. J Pathol, 2008. 214(2): p. 231–41.

14. Nixon, D.E. and A.L. Landay, Biomarkers of immune dysfunction in HIV. Current Opinion in HIV and AIDS, 2010. 5(6): p. 498–503.

15. Boasso, A. and G.M. Shearer, Chronic innate immune activation as a cause of HIV-1 immunopathogenesis. Clinical Immunology, 2008. 126(3): p. 235–242.

16. Chang, J.J. and M. Altfeld, Innate Immune Activation in Primary HIV-1 Infection. The Journal of Infectious Diseases, 2010. 202(S2): p. S297–S301.

17. Deeks, S.G., C.M. Kitchen, L. Liu, H. Guo, R. Gascon, A.B. Narvaez, P. Hunt, J.N. Martin, J.O. Kahn, J. Levy, M.S. McGrath, and F.M. Hecht, Immune activation set point during early HIV infection predicts subsequent CD4+ T-cell changes independent of viral load. Blood, 2004. 104(4): p. 942–7.

18. Douek, D., HIV disease progression: immune activation, microbes, and a leaky gut. Top HIV Med, 2007. 15(4): p. 114–7.

19. Hazenberg, M.D., S.A. Otto, B.H. van Benthem, M.T. Roos, R.A. Coutinho, J.M. Lange, D. Hamann, M. Prins, and F. Miedema, Persistent immune activation in HIV-1 infection is associated with progression to AIDS. AIDS, 2003. 17(13): p. 1881–8.

20. Sousa, A.E., J. Carneiro, M. Meier-Schellersheim, Z. Grossman, and R.M. Victorino, CD4 T cell depletion is linked directly to immune activation in the pathogenesis of HIV-1 and HIV-2 but only indirectly to the viral load. J Immunol, 2002. 169(6): p. 3400–6.

21. Feldman, S.B., M.C. Milone, P. Kloser, and P. Fitzgerald-Bocarsly, Functional deficiencies in two distinct interferon alpha-producing cell populations in peripheral blood mononuclear cells from human immunodeficiency virus seropositive patients. J Leukoc Biol, 1995. 57(2): p. 214–20.

22. Ronnblom, L., U. Ramstedt, and G.V. Alm, Properties of human natural interferon-producing cells stimulated by tumor cell lines. Eur J Immunol, 1983. 13(6): p. 471–6.

23. Fitzgerald-Bocarsly, P. and E.S. Jacobs, Plasmacytoid dendritic cells in HIV infection: striking a delicate balance. Journal of Leukocyte Biology, 2010. 87(4): p. 609–620.

24. Kadowaki, N., S. Antonenko, J.Y. Lau, and Y.J. Liu, Natural interferon alpha/beta-producing cells link innate and adaptive immunity. J Exp Med, 2000. 192(2): p. 219–26.

25. Patterson, S., A. Rae, N. Hockey, J. Gilmour, and F. Gotch, Plasmacytoid dendritic cells are highly susceptible to human immunodeficiency virus type 1 infection and release infectious virus. J Virol, 2001. 75(14): p. 6710–3.

26. Feldman, S., D. Stein, S. Amrute, T. Denny, Z. Garcia, P. Kloser, Y. Sun, N. Megjugorac, and P. Fitzgerald-Bocarsly, Decreased interferon-alpha production in HIV-infected patients correlates with numerical and functional deficiencies in circulating type 2 dendritic cell precursors. Clin Immunol, 2001. 101(2): p. 201–10.

27. Pacanowski, J., S. Kahi, M. Baillet, P. Lebon, C. Deveau, C. Goujard, L. Meyer, E. Oksenhendler, M. Sinet, and A. Hosmalin, Reduced blood CD123+ (lymphoid) and CD11c+ (myeloid) dendritic cell numbers in primary HIV-1 infection. Blood, 2001. 98(10): p. 3016–21.

28. Soumelis, V., I. Scott, F. Gheyas, D. Bouhour, G. Cozon, L. Cotte, L. Huang, J.A. Levy, and Y.J. Liu, Depletion of circulating natural type 1 interferon-producing cells in HIV-infected AIDS patients. Blood, 2001. 98(4): p. 906–12.

29. Donaghy, H., A. Pozniak, B. Gazzard, N. Qazi, J. Gilmour, F. Gotch, and S. Patterson, Loss of blood CD11c(+) myeloid and CD11c(-) plasmacytoid dendritic cells in patients with HIV-1 infection correlates with HIV-1 RNA virus load. Blood, 2001. 98(8): p. 2574–6.

30. Kamga, I., S. Kahi, L. Develioglu, M. Lichtner, C. Maranon, C. Deveau, L. Meyer, C. Goujard, P. Lebon, M. Sinet, and A. Hosmalin, Type I interferon production is profoundly and transiently impaired in primary HIV-1 infection. J Infect Dis, 2005. 192(2): p. 303–10.

31. Hosmalin, A. and P. Lebon, Type I interferon production in HIV-infected patients. Journal of Leukocyte Biology, 2006. 80(5): p. 984–993.

32. Cavaleiro, R., A.P. Baptista, R.S. Soares, R. Tendeiro, R.B. Foxall, P. Gomes, R.M. Victorino, and A.E. Sousa, Major depletion of plasmacytoid dendritic cells in HIV-2 infection, an attenuated form of HIV disease. PLoS Pathog, 2009. 5(11): p. e1000667.

33. Chehimi, J., D.E. Campbell, L. Azzoni, D. Bacheller, E. Papasavvas, G. Jerandi, K. Mounzer, J. Kostman, G. Trinchieri, and L.J. Montaner, Persistent decreases in blood plasmacytoid dendritic cell number and function despite effective highly active antiretroviral therapy and increased blood myeloid dendritic cells in HIV-infected individuals. J Immunol, 2002. 168(9): p. 4796–801.

34. Donaghy, H., B. Gazzard, F. Gotch, and S. Patterson, Dysfunction and infection of freshly isolated blood myeloid and plasmacytoid dendritic cells in patients infected with HIV-1. Blood, 2003. 101(11): p. 4505–11.

35. Martinson, J.A., A. Roman-Gonzalez, A.R. Tenorio, C.J. Montoya, C.N. Gichinga, M.T. Rugeles, M. Tomai, A.M. Krieg, S. Ghanekar, L.L. Baum, and A.L. Landay, Dendritic cells from HIV-1 infected individuals are less responsive to toll-like receptor (TLR) ligands. Cell Immunol, 2007. 250(1-2): p. 75–84.

36. Tilton, J.C., M.M. Manion, M.R. Luskin, A.J. Johnson, A.A. Patamawenu, C.W. Hallahan, N.A. Cogliano-Shutta, J.M. Mican, R.T. Davey, Jr., S. Kottilil, J.D. Lifson, J.A. Metcalf, R.A. Lempicki, and M. Connors, Human immunodeficiency virus viremia induces plasmacytoid dendritic cell activation in vivo and diminished alpha interferon production in vitro. J Virol, 2008. 82(8): p. 3997–4006.

37. Conry, S.J., K.A. Milkovich, N.L. Yonkers, B. Rodriguez, H.B. Bernstein, R. Asaad, F.P. Heinzel, M. Tary-Lehmann, M.M. Lederman, and D.D. Anthony, Impaired plasmacytoid dendritic cell (PDC)-NK cell activity in viremic human immunodeficiency virus infection attributable to impairments in both PDC and NK cell function. J Virol, 2009. 83(21): p. 11175–87.

38. Reitano, K.N., S. Kottilil, C.M. Gille, X. Zhang, M. Yan, M.A. O’Shea, G. Roby, C.W. Hallahan, J. Yang, R.A. Lempicki, J. Arthos, and A.S. Fauci, Defective plasmacytoid dendritic cell-NK cell cross-talk in HIV infection. AIDS Res Hum Retroviruses, 2009. 25(10): p. 1029–37.

39. Brown, K.N., A. Trichel, and S.M. Barratt-Boyes, Parallel loss of myeloid and plasmacytoid dendritic cells from blood and lymphoid tissue in simian AIDS. J Immunol, 2007. 178(11): p. 6958–67.

40. Jenabian, M.A., M. El-Far, K. Vyboh, I. Kema, C.T. Costiniuk, R. Thomas, J.G. Baril, R. LeBlanc, C. Kanagaratham, D. Radzioch, O. Allam, A. Ahmad, B. Lebouche, C. Tremblay, P. Ancuta, J.P. Routy, i. for the Montreal Primary, and G. Slow Progressor Study, Immunosuppressive Tryptophan Catabolism and Gut Mucosal Dysfunction Following Early HIV Infection. J Infect Dis, 2015.

41. Colonna, M., Toll-like receptors and IFN-alpha: partners in autoimmunity. J Clin Invest, 2006. 116(9): p. 2319–22.

42. Boasso, A., J.P. Herbeuval, A.W. Hardy, S.A. Anderson, M.J. Dolan, D. Fuchs, and G.M. Shearer, HIV inhibits CD4+ T-cell proliferation by inducing indoleamine 2,3-dioxygenase in plasmacytoid dendritic cells. Blood, 2007. 109(8): p. 3351–9.

43. Manches, O., D. Munn, A. Fallahi, J. Lifson, L. Chaperot, J. Plumas, and N. Bhardwaj, HIV-activated human plasmacytoid DCs induce Tregs through an indoleamine 2,3-dioxygenase-dependent mechanism. J Clin Invest, 2008. 118(10): p. 3431–9.

44. Effros, R.B., C.V. Fletcher, K. Gebo, J.B. Halter, W.R. Hazzard, F.M. Horne, R.E. Huebner, E.N. Janoff, A.C. Justice, D. Kuritzkes, S.G. Nayfield, S.F. Plaeger, K.E. Schmader, J.R. Ashworth, C. Campanelli, C.P. Clayton, B. Rada, N.F. Woolard, and K.P. High, Aging and infectious diseases: workshop on HIV infection and aging: what is known and future research directions. Clin Infect Dis, 2008. 47(4): p. 542–53.

45. Klatt, N.R., N. Chomont, D.C. Douek, and S.G. Deeks, Immune activation and HIV persistence: implications for curative approaches to HIV infection. Immunol Rev, 2013. 254(1): p. 326–42.

46. Keir, M.E., G.J. Freeman, and A.H. Sharpe, PD-1 regulates self-reactive CD8+ T cell responses to antigen in lymph nodes and tissues. J Immunol, 2007. 179(8): p. 5064–70.

47. Keir, M.E., L.M. Francisco, and A.H. Sharpe, PD-1 and its ligands in T-cell immunity. Curr Opin Immunol, 2007. 19(3): p. 309–14.

48. Wherry, E.J., T cell exhaustion. Nat Immunol, 2011. 12(6): p. 492–9.

49. Freeman, G.J., E.J. Wherry, R. Ahmed, and A.H. Sharpe, Reinvigorating exhausted HIV-specific T cells via PD-1-PD-1 ligand blockade. J Exp Med, 2006. 203(10): p. 2223–7.

50. Khaitan, A. and D. Unutmaz, Revisiting immune exhaustion during HIV infection. Curr HIV/AIDS Rep, 2011. 8(1): p. 4–11.

51. Muthumani, K., D.J. Shedlock, D.K. Choo, P. Fagone, O.U. Kawalekar, J. Goodman, C.B. Bian, A.A. Ramanathan, P. Atman, P. Tebas, M.A. Chattergoon, A.Y. Choo, and D.B. Weiner, HIV-mediated phosphatidylinositol 3-kinase/serine-threonine kinase activation in APCs leads to programmed death-1 ligand upregulation and suppression of HIV-specific CD8 T cells. J Immunol, 2011. 187(6): p. 2932–43.

52. Keir, M., L. Francisco, and A. Sharpe, PD-1 and its ligands in T-cell immunity. Current Opinion in Immunology, 2007. 19(3): p. 309–314.

53. Jarvis, L.M., Releasing the Immune Brakes. Chemical & Engineering News, 2015. 93(14): p. 10-+.

54. Baksh, K. and J. Weber, Immune Checkpoint Protein Inhibition for Cancer: Preclinical Justification for CTLA-4 and PD-1 Blockade and New Combinations. Semin Oncol, 2015. 42(3): p. 363–377.

55. Drake, C.G., E. Jaffee, and D.M. Pardoll, Mechanisms of immune evasion by tumors. Adv Immunol, 2006. 90: p. 51–81.

56. Mizoguchi, H., J.J. O’Shea, D.L. Longo, C.M. Loeffler, D.W. McVicar, and A.C. Ochoa, Alterations in signal transduction molecules in T lymphocytes from tumor-bearing mice. Science, 1992. 258(5089): p. 1795–8.

57. Dong, H., S.E. Strome, D.R. Salomao, H. Tamura, F. Hirano, D.B. Flies, P.C. Roche, J. Lu, G. Zhu, K. Tamada, V.A. Lennon, E. Celis, and L. Chen, Tumor-associated B7-H1 promotes T-cell apoptosis: a potential mechanism of immune evasion. Nat Med, 2002. 8(8): p. 793–800.

58. Meier, A., A. Bagchi, H.K. Sidhu, G. Alter, T.J. Suscovich, D.G. Kavanagh, H. Streeck, M.A. Brockman, S. LeGall, J. Hellman, and M. Altfeld, Upregulation of PD-L1 on monocytes and dendritic cells by HIV-1 derived TLR ligands. AIDS, 2008. 22(5): p. 655–8.

59. Piguet, V., S.M. Caucheteux, M. Iannetta, and A. Hosmalin, Altered antigen-presenting cells during HIV-1 infection. Curr Opin HIV AIDS, 2014. 9(5): p. 478–84.

60. Jin, X., D.E. Bauer, S.E. Tuttleton, S. Lewin, A. Gettie, J. Blanchard, C.E. Irwin, J.T. Safrit, J. Mittler, L. Weinberger, L.G. Kostrikis, L. Zhang, A.S. Perelson, and D.D. Ho, Dramatic rise in plasma viremia after CD8(+) T cell depletion in simian immunodeficiency virus-infected macaques. J Exp Med, 1999. 189(6): p. 991–8.

61. Zuniga, E.I., D.B. McGavern, J.L. Pruneda-Paz, C. Teng, and M.B. Oldstone, Bone marrow plasmacytoid dendritic cells can differentiate into myeloid dendritic cells upon virus infection. Nat Immunol, 2004. 5(12): p. 1227–34.

62. Ito, T., Y.J. Liu, and N. Kadowaki, Functional diversity and plasticity of human dendritic cell subsets. Int J Hematol, 2005. 81(3): p. 188–96.

63. Olweus, J., A. BitMansour, R. Warnke, P.A. Thompson, J. Carballido, L.J. Picker, and F. Lund-Johansen, Dendritic cell ontogeny: a human dendritic cell lineage of myeloid origin. Proc Natl Acad Sci U S A, 1997. 94(23): p. 12551–6.

64. Corcoran, L., I. Ferrero, D. Vremec, K. Lucas, J. Waithman, M. O’Keeffe, L. Wu, A. Wilson, and K. Shortman, The lymphoid past of mouse plasmacytoid cells and thymic dendritic cells. J Immunol, 2003. 170(10): p. 4926–32.

65. Dai, J., N.J. Megjugorac, S.B. Amrute, and P. Fitzgerald-Bocarsly, Regulation of IFN regulatory factor-7 and IFN-alpha production by enveloped virus and lipopolysaccharide in human plasmacytoid dendritic cells. J Immunol, 2004. 173(3): p. 1535–48.

66. Lederman, M.M., L. Calabrese, N.T. Funderburg, B. Clagett, K. Medvik, H. Bonilla, B. Gripshover, R.A. Salata, A. Taege, M. Lisgaris, G.A. McComsey, E. Kirchner, J. Baum, C. Shive, R. Asaad, R.C. Kalayjian, S.F. Sieg, and B. Rodriguez, Immunologic failure despite suppressive antiretroviral therapy is related to activation and turnover of memory CD4 cells. J Infect Dis, 2011. 204(8): p. 1217–26.

67. Hunt, P.W., J.N. Martin, E. Sinclair, B. Bredt, E. Hagos, H. Lampiris, and S.G. Deeks, T cell activation is associated with lower CD4+ T cell gains in human immunodeficiency virus-infected patients with sustained viral suppression during antiretroviral therapy. J Infect Dis, 2003. 187(10): p. 1534–43.

68. Baroncelli, S., C.M. Galluzzo, M.F. Pirillo, M.G. Mancini, L.E. Weimer, M. Andreotti, R. Amici, S. Vella, M. Giuliano, and L. Palmisano, Microbial translocation is associated with residual viral replication in HAART-treated HIV+ subjects with <50copies/ml HIV-1 RNA. J Clin Virol, 2009. 46(4): p. 367–70.

69. Brenchley, J.M., D.A. Price, T.W. Schacker, T.E. Asher, G. Silvestri, S. Rao, Z. Kazzaz, E. Bornstein, O. Lambotte, D. Altmann, B.R. Blazar, B. Rodriguez, L. Teixeira-Johnson, A. Landay, J.N. Martin, F.M. Hecht, L.J. Picker, M.M. Lederman, S.G. Deeks, and D.C. Douek, Microbial translocation is a cause of systemic immune activation in chronic HIV infection. Nat Med, 2006. 12(12): p. 1365–71.

70. Sachdeva, M., M.A. Fischl, R. Pahwa, N. Sachdeva, and S. Pahwa, Immune exhaustion occurs concomitantly with immune activation and decrease in regulatory T cells in viremic chronically HIV-1-infected patients. J Acquir Immune Defic Syndr, 2010. 54(5): p. 447–54.

71. Hunt, P.W., Role of immune activation in HIV pathogenesis. Curr HIV/AIDS Rep, 2007. 4(1): p. 42–7.

72. Hunt, Peter W., J. Brenchley, E. Sinclair, Joseph M. McCune, M. Roland, K. Page-Shafer, P. Hsue, B. Emu, M. Krone, H. Lampiris, D. Douek, Jeffrey N. Martin, and Steven G. Deeks, Relationship between T Cell Activation and CD4+T Cell Count in HIV-Seropositive Individuals with Undetectable Plasma HIV RNA Levels in the Absence of Therapy. The Journal of Infectious Diseases, 2008. 197(1): p. 126–133.

73. Marchetti, G., G.M. Bellistri, E. Borghi, C. Tincati, S. Ferramosca, M. La Francesca, G. Morace, A. Gori, and A.D. Monforte, Microbial translocation is associated with sustained failure in CD4+ T-cell reconstitution in HIV-infected patients on long-term highly active antiretroviral therapy. AIDS, 2008. 22(15): p. 2035–8.

74. Marchetti, G., A. Cozzi-Lepri, E. Merlini, G.M. Bellistri, A. Castagna, M. Galli, G. Verucchi, A. Antinori, A. Costantini, A. Giacometti, A. di Caro, and A. D’Arminio Monforte, Microbial translocation predicts disease progression of HIV-infected antiretroviral-naive patients with high CD4+ cell count. AIDS, 2011. 25(11): p. 1385–94.

75. Gori, A., C. Tincati, G. Rizzardini, C. Torti, T. Quirino, M. Haarman, K. Ben Amor, J. van Schaik, A. Vriesema, J. Knol, G. Marchetti, G. Welling, and M. Clerici, Early Impairment of Gut Function and Gut Flora Supporting a Role for Alteration of Gastrointestinal Mucosa in Human Immunodeficiency Virus Pathogenesis. Journal of Clinical Microbiology, 2007. 46(2): p. 757–758.

76. Hunt, P.W., J. Brenchley, E. Sinclair, J.M. McCune, M. Roland, K. Page-Shafer, P. Hsue, B. Emu, M. Krone, H. Lampiris, D. Douek, J.N. Martin, and S.G. Deeks, Relationship between T cell activation and CD4+ T cell count in HIV-seropositive individuals with undetectable plasma HIV RNA levels in the absence of therapy. J Infect Dis, 2008. 197(1): p. 126–33.

77. Jiang, W., M.M. Lederman, P. Hunt, S.F. Sieg, K. Haley, B. Rodriguez, A. Landay, J. Martin, E. Sinclair, A.I. Asher, S.G. Deeks, D.C. Douek, and J.M. Brenchley, Plasma levels of bacterial DNA correlate with immune activation and the magnitude of immune restoration in persons with antiretroviral-treated HIV infection. J Infect Dis, 2009. 199(8): p. 1177–85.

78. Bukh, A.R., J. Melchjorsen, R. Offersen, J.M. Jensen, L. Toft, H. Stovring, L. Ostergaard, M. Tolstrup, and O.S. Sogaard, Endotoxemia is associated with altered innate and adaptive immune responses in untreated HIV-1 infected individuals. PLoS One, 2011. 6(6): p. e21275.

79. Lee, P.I., E.J. Ciccone, S.W. Read, A. Asher, R. Pitts, D.C. Douek, J.M. Brenchley, and I. Sereti, Evidence for translocation of microbial products in patients with idiopathic CD4 lymphocytopenia. J Infect Dis, 2009. 199(11): p. 1664–70.

80. Troseid, M., A. Sonnerborg, and P. Nowak, High mobility group box protein-1 in HIV-1 infection. Curr HIV Res, 2011. 9(1): p. 6–10.

81. Schmittgen, T.D. and K.J. Livak, Analyzing real-time PCR data by the comparative C(T) method. Nat Protoc, 2008. 3(6): p. 1101–8.

82. Dzionek, A., A. Fuchs, P. Schmidt, S. Cremer, M. Zysk, S. Miltenyi, D.W. Buck, and J. Schmitz, BDCA-2, BDCA-3, and BDCA-4: three markers for distinct subsets of dendritic cells in human peripheral blood. J Immunol, 2000. 165(11): p. 6037–46.

83. Hemmi, H., T. Kaisho, O. Takeuchi, S. Sato, H. Sanjo, K. Hoshino, T. Horiuchi, H. Tomizawa, K. Takeda, and S. Akira, Small anti-viral compounds activate immune cells via the TLR7 MyD88-dependent signaling pathway. Nat Immunol, 2002. 3(2): p. 196–200.

84. Krug, A., A.R. French, W. Barchet, J.A. Fischer, A. Dzionek, J.T. Pingel, M.M. Orihuela, S. Akira, W.M. Yokoyama, and M. Colonna, TLR9-dependent recognition of MCMV by IPC and DC generates coordinated cytokine responses that activate antiviral NK cell function. Immunity, 2004. 21(1): p. 107–19.

85. Lund, J., A. Sato, S. Akira, R. Medzhitov, and A. Iwasaki, Toll-like receptor 9-mediated recognition of Herpes simplex virus-2 by plasmacytoid dendritic cells. J Exp Med, 2003. 198(3): p. 513–20.

86. Beignon, A.S., Endocytosis of HIV-1 activates plasmacytoid dendritic cells via Toll-like receptor-viral RNA interactions. Journal of Clinical Investigation, 2005. 115(11): p. 3265–3275.

87. Piccioli, D., C. Sammicheli, S. Tavarini, S. Nuti, E. Frigimelica, A.G. Manetti, A. Nuccitelli, S. Aprea, S. Valentini, E. Borgogni, A. Wack, and N.M. Valiante, Human plasmacytoid dendritic cells are unresponsive to bacterial stimulation and require a novel type of cooperation with myeloid dendritic cells for maturation. Blood, 2009. 113(18): p. 4232–9.

88. Hernandez, J.C., M. Stevenson, E. Latz, and S. Urcuqui-Inchima, HIV type 1 infection up-regulates TLR2 and TLR4 expression and function in vivo and in vitro. AIDS Res Hum Retroviruses, 2012. 28(10): p. 1313–28.

89. Hernandez, J.C., J. Arteaga, S. Paul, A. Kumar, E. Latz, and S. Urcuqui-Inchima, Up-regulation of TLR2 and TLR4 in dendritic cells in response to HIV type 1 and coinfection with opportunistic pathogens. AIDS Res Hum Retroviruses, 2011. 27(10): p. 1099–109.

90. Asselin-Paturel, C. and G. Trinchieri, Production of type I interferons: plasmacytoid dendritic cells and beyond. J Exp Med, 2005. 202(4): p. 461–5.

91. Zanoni, I. and F. Granucci, Differences in lipopolysaccharide-induced signaling between conventional dendritic cells and macrophages. Immunobiology, 2010. 215(9-10): p. 709–12.

92. Beutler, B., Z. Jiang, P. Georgel, K. Crozat, B. Croker, S. Rutschmann, X. Du, and K. Hoebe, Genetic analysis of host resistance: Toll-like receptor signaling and immunity at large. Annu Rev Immunol, 2006. 24: p. 353–89.

93. Zanoni, I., R. Ostuni, G. Capuano, M. Collini, M. Caccia, A.E. Ronchi, M. Rocchetti, F. Mingozzi, M. Foti, G. Chirico, B. Costa, A. Zaza, P. Ricciardi-Castagnoli, and F. Granucci, CD14 regulates the dendritic cell life cycle after LPS exposure through NFAT activation. Nature, 2009. 460(7252): p. 264–8.

94. Zanoni, I., C. Bodio, A. Broggi, R. Ostuni, M. Caccia, M. Collini, A. Venkatesh, R. Spreafico, G. Capuano, and F. Granucci, Similarities and differences of innate immune responses elicited by smooth and rough LPS. Immunol Lett, 2012. 142(1-2): p. 41–7.

95. Zanoni, I., R. Ostuni, L.R. Marek, S. Barresi, R. Barbalat, G.M. Barton, F. Granucci, and J.C. Kagan, CD14 controls the LPS-induced endocytosis of Toll-like receptor 4. Cell, 2011. 147(4): p. 868–80.

96. Jiang, Z., P. Georgel, X. Du, L. Shamel, S. Sovath, S. Mudd, M. Huber, C. Kalis, S. Keck, C. Galanos, M. Freudenberg, and B. Beutler, CD14 is required for MyD88-independent LPS signaling. Nat Immunol, 2005. 6(6): p. 565–70.

97. Alcami, J., T. Lain de Lera, L. Folgueira, M.A. Pedraza, J.M. Jacque, F. Bachelerie, A.R. Noriega, R.T. Hay, D. Harrich, R.B. Gaynor, and, et al., Absolute dependence on kappa B responsive elements for initiation and Tat-mediated amplification of HIV transcription in blood CD4 T lymphocytes. EMBO J, 1995. 14(7): p. 1552–60.

98. Liou, H.C. and D. Baltimore, Regulation of the NF-kappa B/rel transcription factor and I kappa B inhibitor system. Curr Opin Cell Biol, 1993. 5(3): p. 477–87.

99. Kawamoto, T., M. Ii, T. Kitazaki, Y. Iizawa, and H. Kimura, TAK-242 selectively suppresses Toll-like receptor 4-signaling mediated by the intracellular domain. Eur J Pharmacol, 2008. 584(1): p. 40–8.

100. Roulston, A., R. Lin, P. Beauparlant, M.A. Wainberg, and J. Hiscott, Regulation of human immunodeficiency virus type 1 and cytokine gene expression in myeloid cells by NF-kappa B/Rel transcription factors. Microbiol Rev, 1995. 59(3): p. 481–505.

101. Ryan, L.K., G. Diamond, S. Amrute, Z. Feng, A. Weinberg, and P. Fitzgerald-Bocarsly, Detection of HBD1 peptide in peripheral blood mononuclear cell subpopulations by intracellular flow cytometry. Peptides, 2003. 24(11): p. 1785–94.

102. Kuller, L.H., R. Tracy, W. Belloso, S. De Wit, F. Drummond, H.C. Lane, B. Ledergerber, J. Lundgren, J. Neuhaus, D. Nixon, N.I. Paton, J.D. Neaton, and I.S.S. Group, Inflammatory and coagulation biomarkers and mortality in patients with HIV infection. PLoS Med, 2008. 5(10): p. e203.

103. Rodger, A.J., Z. Fox, J.D. Lundgren, L.H. Kuller, C. Boesecke, D. Gey, A. Skoutelis, M.B. Goetz, A.N. Phillips, and I.S.f.M.o.A.T.S. Group, Activation and coagulation biomarkers are independent predictors of the development of opportunistic disease in patients with HIV infection. J Infect Dis, 2009. 200(6): p. 973–83.

104. Hosmalin, A. and P. Lebon, Type I interferon production in HIV-infected patients. J Leukoc Biol, 2006. 80(5): p. 984–93.

105. Yoneyama, H., K. Matsuno, E. Toda, T. Nishiwaki, N. Matsuo, A. Nakano, S. Narumi, B. Lu, C. Gerard, S. Ishikawa, and K. Matsushima, Plasmacytoid DCs help lymph node DCs to induce anti-HSV CTLs. J Exp Med, 2005. 202(3): p. 425–35.

106. Cella, M., D. Jarrossay, F. Facchetti, O. Alebardi, H. Nakajima, A. Lanzavecchia, and M. Colonna, Plasmacytoid monocytes migrate to inflamed lymph nodes and produce large amounts of type I interferon. Nat Med, 1999. 5(8): p. 919–23.

107. Megjugorac, N.J., Virally stimulated plasmacytoid dendritic cells produce chemokines and induce migration of T and NK cells. Journal of Leukocyte Biology, 2003. 75(3): p. 504–514.

108. Fonteneau, J.F., M. Larsson, A.S. Beignon, K. McKenna, I. Dasilva, A. Amara, Y.J. Liu, J.D. Lifson, D.R. Littman, and N. Bhardwaj, Human immunodeficiency virus type 1 activates plasmacytoid dendritic cells and concomitantly induces the bystander maturation of myeloid dendritic cells. J Virol, 2004. 78(10): p. 5223–32.

109. Herbeuval, J.P., J. Nilsson, A. Boasso, A.W. Hardy, M.J. Kruhlak, S.A. Anderson, M.J. Dolan, M. Dy, J. Andersson, and G.M. Shearer, Differential expression of IFN-alpha and TRAIL/DR5 in lymphoid tissue of progressor versus nonprogressor HIV-1-infected patients. Proc Natl Acad Sci U S A, 2006. 103(18): p. 7000–5.

110. Lester, R.T., X.D. Yao, T.B. Ball, L.R. McKinnon, R. Kaul, C. Wachihi, W. Jaoko, F.A. Plummer, and K.L. Rosenthal, Toll-like receptor expression and responsiveness are increased in viraemic HIV-1 infection. AIDS, 2008. 22(6): p. 685–94.

111. Bafica, A., C.A. Scanga, M. Schito, D. Chaussabel, and A. Sher, Influence of coinfecting pathogens on HIV expression: evidence for a role of Toll-like receptors. J Immunol, 2004. 172(12): p. 7229–34.

112. Beignon, A.S., K. McKenna, M. Skoberne, O. Manches, I. DaSilva, D.G. Kavanagh, M. Larsson, R.J. Gorelick, J.D. Lifson, and N. Bhardwaj, Endocytosis of HIV-1 activates plasmacytoid dendritic cells via Toll-like receptor-viral RNA interactions. J Clin Invest, 2005. 115(11): p. 3265–75.

113. Sandler, N.G., H. Wand, A. Roque, M. Law, M.C. Nason, D.E. Nixon, C. Pedersen, K. Ruxrungtham, S.R. Lewin, S. Emery, J.D. Neaton, J.M. Brenchley, S.G. Deeks, I. Sereti, and D.C. Douek, Plasma levels of soluble CD14 independently predict mortality in HIV infection. J Infect Dis, 2011. 203(6): p. 780–90.

114. Mavigner, M., M. Cazabat, M. Dubois, F.E. L’Faqihi, M. Requena, C. Pasquier, P. Klopp, J. Amar, L. Alric, K. Barange, J.P. Vinel, B. Marchou, P. Massip, J. Izopet, and P. Delobel, Altered CD4+ T cell homing to the gut impairs mucosal immune reconstitution in treated HIV-infected individuals. J Clin Invest, 2012. 122(1): p. 62–9.

115. Merlini, E., F. Bai, G.M. Bellistri, C. Tincati, A. d’Arminio Monforte, and G. Marchetti, Evidence for polymicrobic flora translocating in peripheral blood of HIV-infected patients with poor immune response to antiretroviral therapy. PLoS One, 2011. 6(4): p. e18580.

116. Papasavvas, E., M. Pistilli, G. Reynolds, R. Bucki, L. Azzoni, J. Chehimi, P.A. Janmey, M.J. DiNubile, J. Ondercin, J.R. Kostman, K.C. Mounzer, and L.J. Montaner, Delayed loss of control of plasma lipopolysaccharide levels after therapy interruption in chronically HIV-1-infected patients. AIDS, 2009. 23(3): p. 369–75.

117. Kramski, M., A.J. Gaeguta, G.F. Lichtfuss, R. Rajasuriar, S.M. Crowe, M.A. French, S.R. Lewin, R.J. Center, and D.F. Purcell, Novel sensitive real-time PCR for quantification of bacterial 16S rRNA genes in plasma of HIV-infected patients as a marker for microbial translocation. J Clin Microbiol, 2011. 49(10): p. 3691–3.

118. Equils, O., E. Faure, L. Thomas, Y. Bulut, S. Trushin, and M. Arditi, Bacterial lipopolysaccharide activates HIV long terminal repeat through Toll-like receptor 4. J Immunol, 2001. 166(4): p. 2342–7.

119. Nazli, A., J.K. Kafka, V.H. Ferreira, V. Anipindi, K. Mueller, B.J. Osborne, S. Dizzell, S. Chauvin, M.F. Mian, M. Ouellet, M.J. Tremblay, K.L. Mossman, A.A. Ashkar, C. Kovacs, D.M. Bowdish, D.P. Snider, R. Kaul, and C. Kaushic, HIV-1 gp120 induces TLR2- and TLR4-mediated innate immune activation in human female genital epithelium. J Immunol, 2013. 191(8): p. 4246–58.

120. Fanning, S.L., T.C. George, D. Feng, S.B. Feldman, N.J. Megjugorac, A.G. Izaguirre, and P. Fitzgerald-Bocarsly, Receptor cross-linking on human plasmacytoid dendritic cells leads to the regulation of IFN-alpha production. J Immunol, 2006. 177(9): p. 5829–39.

121. Landmann, R., H.P. Knopf, S. Link, S. Sansano, R. Schumann, and W. Zimmerli, Human monocyte CD14 is upregulated by lipopolysaccharide. Infect Immun, 1996. 64(5): p. 1762–9.

122. Velu, V., R.D. Shetty, M. Larsson, and E.M. Shankar, Role of PD-1 co-inhibitory pathway in HIV infection and potential therapeutic options. Retrovirology, 2015. 12: p. 14.

123. Yamamoto, T., D.A. Price, J.P. Casazza, G. Ferrari, M. Nason, P.K. Chattopadhyay, M. Roederer, E. Gostick, P.D. Katsikis, D.C. Douek, R. Haubrich, C. Petrovas, and R.A. Koup, Surface expression patterns of negative regulatory molecules identify determinants of virus-specific CD8+ T-cell exhaustion in HIV infection. Blood, 2011. 117(18): p. 4805–4815.

124. Planes, R., L. BenMohamed, K. Leghmari, P. Delobel, J. Izopet, and E. Bahraoui, HIV-1 Tat protein induces PD-L1 (B7-H1) expression on dendritic cells through tumor necrosis factor alpha- and toll-like receptor 4-mediated mechanisms. J Virol, 2014. 88(12): p. 6672–89.

125. Ben Haij, N., K. Leghmari, R. Planes, N. Thieblemont, and E. Bahraoui, HIV-1 Tat protein binds to TLR4-MD2 and signals to induce TNF-alpha and IL-10. Retrovirology, 2013. 10: p. 123.

126. Fitzgerald-Bocarsly, P. and E.S. Jacobs, Plasmacytoid dendritic cells in HIV infection: striking a delicate balance. J Leukoc Biol, 2010. 87(4): p. 609–20.

